# The balance between toxic versus nontoxic microRNAs determines platinum sensitivity in ovarian cancer

**DOI:** 10.1101/2021.01.24.427815

**Authors:** Monal Patel, Yinu Wang, Elizabeth T. Bartom, Rohin Dhir, Kenneth P. Nephew, Daniela Matei, Andrea E. Murmann, Ernst Lengyel, Marcus E. Peter

## Abstract

Numerous micro(mi)RNAs (short noncoding RNAs that negatively regulate gene expression) have been linked to platinum (Pt) sensitivity and resistance in ovarian cancer (OC). miRNA activity occurs when the guide strand of the miRNA, with its seed sequence (pos. 2-7/8), is loaded into the RNA induced silencing complex (RISC) and targets complementary short seed matches in the 3’ untranslated region of mRNAs. Toxic seeds, targeting genes critical for cancer cell survival, have been found in tumor suppressive miRNAs. Many si- and shRNAs can also kill cancer cells via toxic seeds, the most toxic carrying G-rich 6mer seed sequences. We now show that treatment of OC cells with Pt leads to an increase in RISC-bound miRNAs carrying toxic 6mer seeds and a decrease in miRNAs with nontoxic seeds. Pt-resistant cells did not exhibit this toxicity shift but retained sensitivity to cell death mediated by siRNAs carrying toxic 6mer seeds. Analysis of RISC-bound miRNAs in OC patients revealed that the ratio between miRNAs with toxic versus miRNAs with nontoxic seeds was predictive of treatment outcome. Application of the 6mer seed toxicity concept to cancer relevant miRNAs provides a new framework for understanding and predicting cancer therapy responses.

## INTRODUCTION

Micro (mi)RNAs are short 19-22 nucleotide (nt) double-stranded (ds) noncoding RNAs that negatively regulate gene expression. The human genome is estimated to code for ~8300 miRNAs (1). Important regulators of differentiation and development, they are deregulated in virtually all human cancers and can function as tumor suppressors or oncogenes (2–4). miRNA activity involves only a very short region of complete complementarity between the ‘seed’, located at positions 2-7/8 of the guide strand. (5, 6) and the ‘seed matches’ predominantly located in the 3’ untranslated region (3’ UTR) of targeted mRNAs (7, 8) resulting in gene silencing (9). To function, mature ds miRNA duplexes are loaded onto Argonaute (Ago) proteins forming the RNA-induced silencing complex (RISC) (10). The active miRNA guide strand incorporates into the RISC (10), while the inactive passenger strand is degraded (11).

Epithelial ovarian cancer (OC) is one of the deadliest gynecological malignancies (12). Despite recent improvements in our understanding of OC biology (13) and the development of new treatments (14), the vast majority of women with advanced stage OC will experience recurrence and succumb to the disease. Most develop resistance to platinum (Pt) chemotherapy, which is part of the standard first line treatment for OC (15). A large body of literature supports the role of aberrantly expressed miRNAs in the disease course of OC, impacting sensitivity to treatment and long-term treatment outcome. At least 87 miRNAs have been reported to either confer acquired therapy resistance, re-sensitize OC cells to treatment, or to correlate with or predict OC treatment outcome (Supplementary Tables S1 and S2). However, there is very little agreement or overlap in those miRNAs identified, raising the question of how this bewildering array of different miRNAs can be linked to therapy resistance in OC.

We previously discovered that a large number of small interfering (si) and short hairpin (sh) RNAs were toxic to all cancer cells independent of their intended target (16). Cells died of simultaneous activation of multiple cell death pathways (16) and could not become resistant to treatment (17). Subsequently, we found that Ago2, a critical component of the RISC, was required for this cell death, and that the toxic si/shRNAs acted like miRNAs by targeting the 3’ UTRs of genes essential for cell survival (18, 19). A 6mer seed present in the guide/antisense strand of the si/shRNA duplex (18) is sufficient for Death Induced by Survival gene Elimination (DISE) (18, 20). The DISE concept was recently confirmed for prostate cancer (20).

In an arrayed screen of all possible 4096 6mer seeds in a neutral backbone 19mer siRNA, we determined that the most toxic 6mer seeds were universally high in G nucleotides at the 5’ end of the seed, largely independent of tissue origin and species ((21) and 6merdb.org). In human cells, we recently identified the seed consensus GGGGGC (targeting the seed match GCCCCC in a miRNA like fashion) as the most toxic seed sequence across three cell lines (19). Toxic 6mer seeds are found in tumor-suppressive miRNAs, including the prototypical tumor suppressive miRNA families, miR-34/449-5p and miR-15/16-5p (21, 22), and we and others have shown that 6mer seed toxicity (6mer Seed Tox) can be developed for cancer treatment (17, 23). When we treated xenografted OC with siRNAs containing toxic seeds, normal tissues were unaffected (17). Various cancer cell lines became more sensitive to 6mer Seed Tox after EMT was induced or their cancer stemness was increased (24). Both EMT and cancer cell stemness are involved in well-documented mechanisms of chemotherapy resistance (25).

After analyzing miRNAs based on the toxicity of their 6mer seed, we now report that, in cancer cell lines both *in vitro* and *in vivo*, and OC patients, the balance of miRNAs containing toxic *versus* nontoxic seeds strongly associates with the outcome of Pt-based chemotherapy. Analyzing the 6mer seed composition of RISC bound miRNAs may predict which patients are Pt resistant at the time of initial diagnostic surgery or may acquire Pt resistance.

## RESULTS

### Both Toxic siRNAs and Carboplatin Exert Toxicity Through RNAi

We decided to use the two highly toxic seeds, GGCAGU (found in miR-34a-5p) and GGGGGC (found in miR-1237-5p), embedded in an siRNA to probe cells. Both siGGCAGU and siGGGGGC were more toxic to Dicer deficient 293T than wt cells (Supplementary Fig. S1A-S1C). We concluded that both siRNAs killed cells through RNAi, since both showed severely reduced toxicity to Ago2 k.o. HeLa cells, yet were highly toxic to wt HeLa cells (Supplementary Fig. S1D). Similarly, these two siRNAs were also more toxic to wt than to Ago2 k.o. 293T cells (Supplementary Fig. S1E). We previously reported that HCT116 cells deficient for either Drosha or Dicer, which are devoid of canonical miRNAs, were hypersensitive to 6mer Seed Tox and cisplatin (18). We now show that Dicer k.o. 293T cells are also much more sensitive to the toxic effects of clinically used carboplatin (Supplementary Fig. S1F) while isogenic Ago2 k.o. 293T cells showed a reduced susceptibility to the toxicity of carboplatin (Supplementary Fig. S1G). These data suggest that the composition of RISC bound miRNAs and RNAi not only determines the activity of toxic siRNAs but, at least in part, the sensitivity of cells to Pt.

### Platinum-Tolerant OC Cells Are Sensitive to 6mer Seed Tox

To determine whether 6mer Seed Tox contributes to the response of OC cells to Pt, we first tested two cell line pairs representing a Pt sensitive (Pt-S) and resistant (Pt-R) phenotype, respectively. A2780R cells were made resistant by culturing A2780 cells for 5 cycles with increasing concentrations of cisplatin (26). PEO1 and PEO4 cells were derived from the same patient at different times after combination chemotherapy with cisplatin, chlorambucil and 5-fluorouracil (27). PEO4 was a later isolate, taken from the patient after her second relapse and hence more resistant to Pt drugs. Treatment of the A2780 cell pair with cisplatin confirmed that the A2780R cells were more resistant to cisplatin than the parental cells (Fig. 1A and B). Interestingly, A2780R cells were at least as sensitive to both siGGCAGU and siGGGGGC as the parental cells (Fig. 1C and D). A similar phenomenon was observed when the PEO1/4 cell pair were treated with carboplatin or toxic siRNAs (Fig. 1E-H). These data suggest that 6mer Seed Tox can kill platinum resistant OC cells.

**Figure 1.**
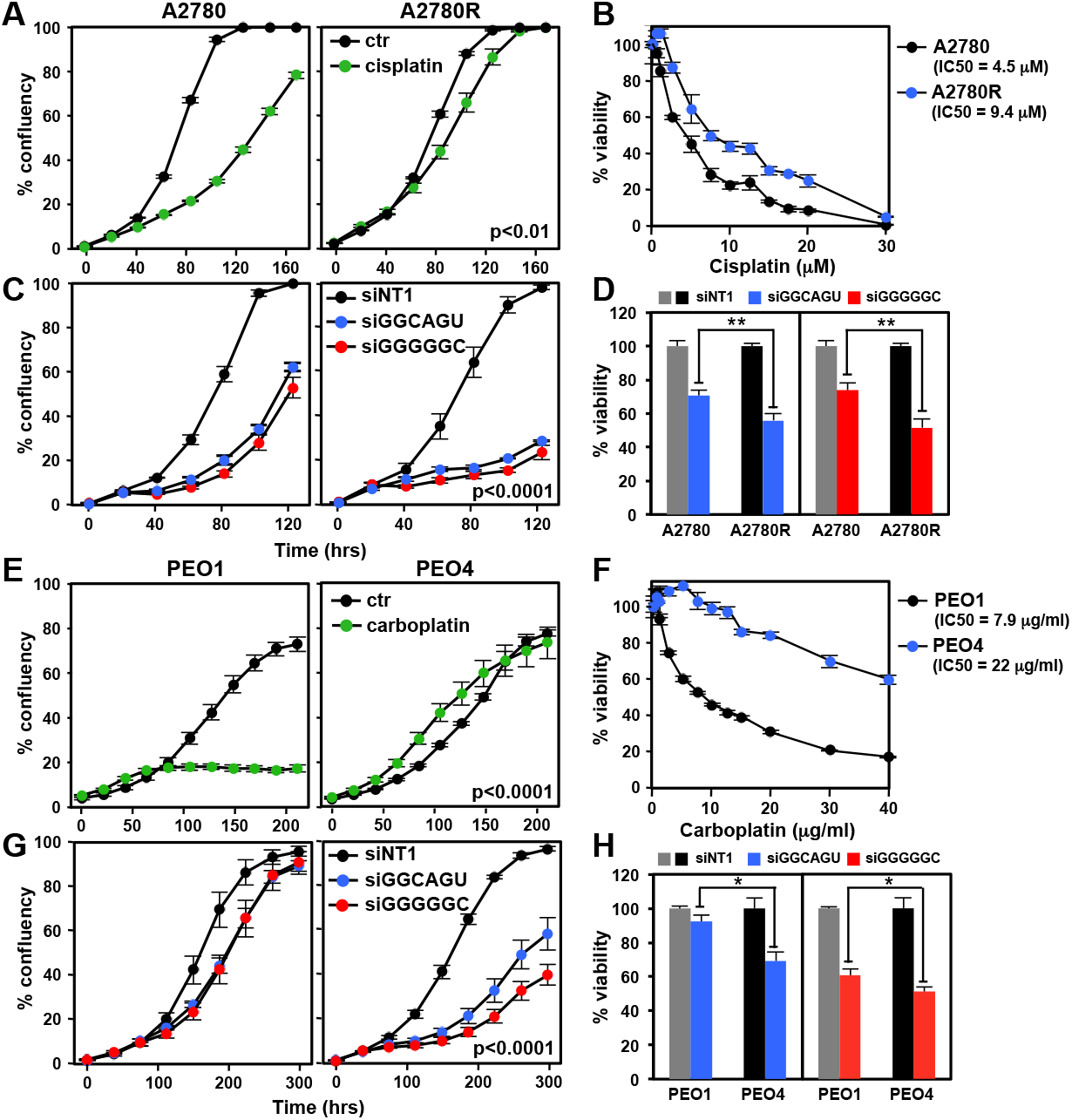
Platinum tolerant Pt-R OC cells are at least as sensitive to toxic 6mer seeds as Pt-S cells. **A,** Change of confluency of A2780 and A2780R cells over time treated with 5 μM Cis. **B,** Dose response curves (96 hrs ATP viability assay) of cells in A. **C,** Change of confluency of A2780 and A2780R cells over time treated with either 10 nM siNT1, siGGCAGU, or siGGGGGC. **D,** Viability of the cells in C after transfection with 10 nM of the siRNAs. **E,** Change of confluency of PEO1 and PEO4 cells over time treated with 1 μg/ml carboplatin. **F,** Dose response curves (96 hrs ATP viability assay) of cells in E. **G,** Change of confluency of PEO1 and PEO4 cells over time treated with either 10 nM siGGCAGU or siGGGGGC. **H,** Viability of the cells in G after transfection with 10 nM of the siRNAs. Each data point (A, B, C, E, F & G) represents mean ± SE of three replicates. Each bar (D, H) represents ± SD of three replicates. P-values were calculated using ANOVA (A, C, E, G), or student’s t-test (D, H). ** p<0.001, * p<0.05.

### Presenting the miRNA Space as a Function of Seed Toxicity - The Seed Tox Graph

Interestingly, many seemingly unrelated miRNAs share seeds of similar toxicity. Therefore, we developed a graphical representation of all miRNAs in cells or tissues at their actual expression levels, plotting the miRNAs according to their seed toxicities (the Seed Tox graph). When comparing miRNA expression in different human tissues, we found that most tissues express predominantly nontoxic seed-containing miRNAs (Supplementary Fig. S2), and only a small fraction of miRNAs had a predicted seed with high toxicity. A few abundant miRNAs are expressed across all tissues (Supplementary Fig. S2). These include the highly toxic miRNAs 22-3p and 24-3p (red), the intermediately toxic miRNAs miR-126-3p, miR-23b-3p, miR-29a-3p (blue), and the nontoxic miRNA members of the let-7 family, miR-125b-5p and miR-21-5p (green).

### Enrichment of miRNAs in the RISC in OC Cells

miRNAs in cancer are almost always studied by quantifying them in the total RNA fraction. However, previous reports demonstrated that most mature miRNAs are not Ago associated (28) and that endogenous miRNAs vary widely in their level of RISC association (29, 30). To assess the 6mer Seed Tox of biologically relevant miRNAs, we pulled down all Ago proteins and their associated miRNAs (18) from two OC cell lines, allowing us to determine the exact positions 2-7 of RISC-bound guide strands. The results were compared to the analysis of total small RNAs in a Seed Tox graph with the x-axis showing the seed toxicity (Supplementary Fig. S3A and S3B). A number of miRNAs were highly enriched in the RISC-bound fraction of both cell lines. These included the toxic miRNAs 22-3p and 24-3p and the nontoxic miRNA miR-125b-5p (Supplementary Fig. S3). In addition, members of the toxic miRNA family miR-15/16, miR-15a/b-5p and miR-424-5p were also highly enriched (Supplementary Fig. S3C). Other miRNAs, most notably the highly abundant members of the let-7-5p family, were reduced in the RISC (Supplementary Fig. S3C). It was also recently reported that many of these miRNAs had a different presence in the RISC than in the total RNA fraction of 293T and A549 cells (asterisks in Supplementary Fig. S3C, (29)). This analysis demonstrates that analyzing RISC bound miRNAs, rather than quantifying total miRNA levels, will more accurately assess functionally relevant miRNAs, which is critical for the analysis of 6mer Seed Tox. We, therefore, used these techniques in the subsequent experiments.

### Changes in RISC Composition Related to Short and Long-Term Treatment with Platinum

Next, we determined the toxicity of RISC bound miRNAs in Pt-S OC cells short-term treated with Pt-based chemotherapy or in Pt-R (long-term treated) cells (Fig. 2). The RISC of A2780 (Pt-S) and A2780R (Pt-R) cells untreated or treated with cisplatin for 72 hrs (a time point before the onset of cell death) was pulled down and the toxicity of all detected miRNAs plotted (Fig. 2A). In short-term cisplatin treated A2780 cells, highly toxic RISC-bound miRNAs (viability <20%), including miR-22/24-3p, were significantly upregulated, while miRNAs with nontoxic seeds (viability >80%) were downregulated (Fig. 2D). The cisplatin induced increase in toxic miRNAs was no longer significant in the Pt-resistant A2780R cells (Fig. 2B). We then compared the RISC content between Pt-S and Pt-R cells (Fig. 2C). The A2780R cells contained fewer toxic miRNAs and more nontoxic miRNAs in the RISC compared to the Pt-S cells (Fig. 2D). A striking difference between the Pt-S and Pt-R cells was the increase in miR-21-5p in the Pt-R cells (Fig. 2C and Supplementary Fig. S4A). Inhibiting miR-21-5p (Supplementary Fig. S4B) rendered Pt-R cells more sensitive to cisplatin (Supplementary Fig. S4C). None of these specific changes were seen when the toxicity of total miRNAs was analyzed (Supplementary Fig. S5) validating this approach of analyzing RISC bound miRNAs. Pt treatment of PEO1 cells, as with treatment of A2780 cells, resulted in a significant increase of RISC bound toxic miRNAs, including 22/24-3p (Fig. 2E). This increase was not found in the Pt-R PEO4 cells (Fig. 2F). Again, the toxic miRNAs were significantly underrepresented in the RISC of the Pt-R PEO4 cells when compared to PEO1 cells (Fig. 2G). Similar changes in the balance between toxic and nontoxic RISC bound miRNAs were therefore found in both OC cell line pairs (Fig. 2D, H). However, nontoxic miRNAs, other than miR-21-5p, were upregulated in the resistant PEO4 cells (green arrows in Fig. 2G). The data suggest that the ratio between miRNAs with toxic versus nontoxic 6mer seeds determines sensitivity to Pt in the OC cell lines.

**Figure 2.**
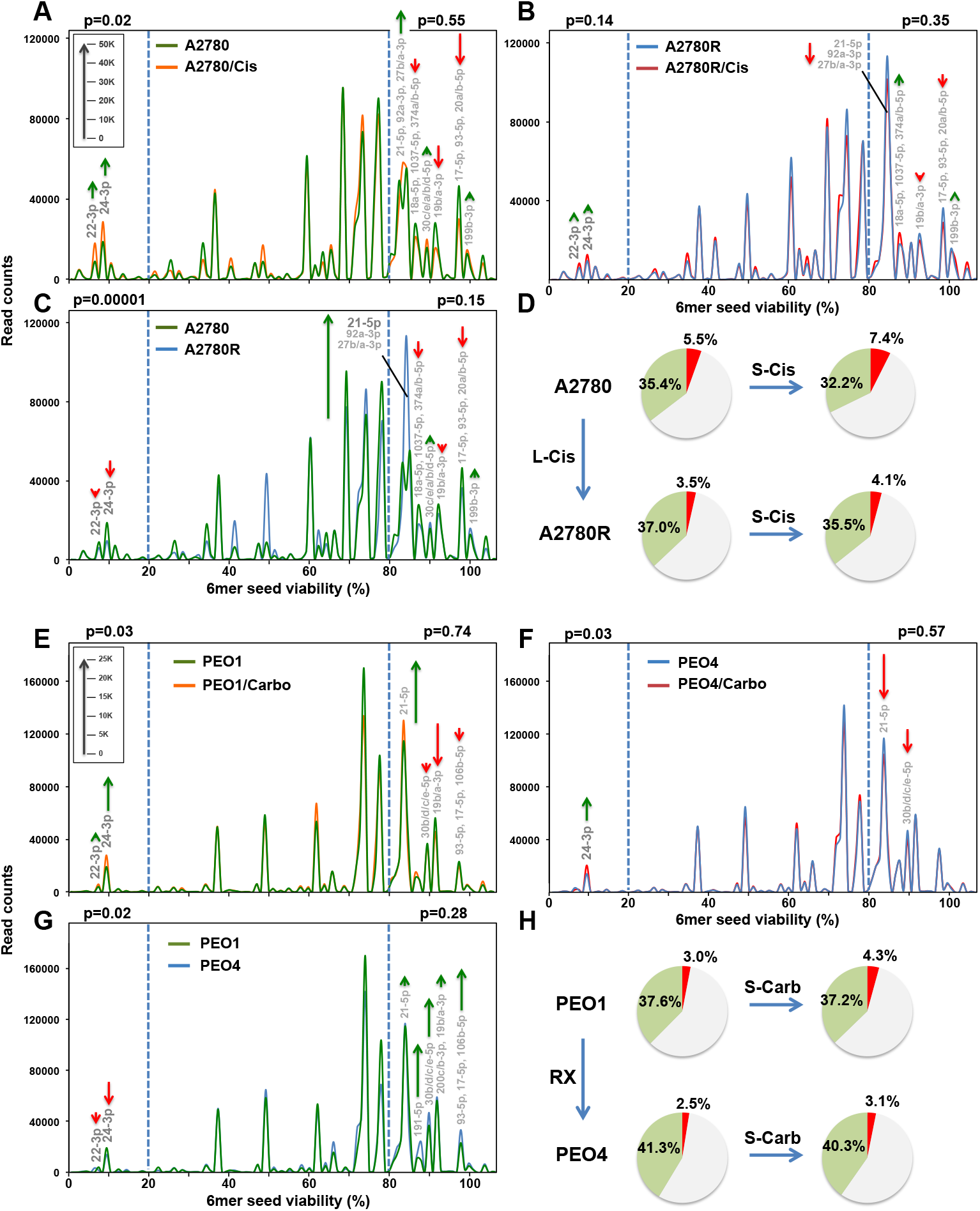
Change in seed toxicity of RISC bound miRNAs in OC cells exposed to platinum. **A** and **B,** RISC-bound sRNA Seed Tox graphs of A2780 (A) or A2780R (B) cells treated with 5 μM Cis or PBS for 72 hrs. Green arrows, net increase, and red arrows, net decrease of >1000 reads in peak in Cis treated cells, respectively. **C,** RISC-bound sRNA Seed Tox graphs of PBS treated A2780 and A2780R cells. Green arrows, net increase, and red arrows, decrease of >1000 reads in peak in A2780R cells, respectively. **D,** RISC composition of A2780 cells or cells made resistant with long-term treatment with cisplatin (A2780R). Shown are pie-charts of short-term (S-Cis) or long-term (L-Cis) treated cells. Reads with a predicted 6mer seed viability of <20% are shown in red, >80% in green, and in between in grey. **E,** RISC-bound sRNA Seed Tox graphs of PEO1 cells treated with 1 μg/ml carboplatin or water for 72 hrs. Green arrows, net increase, and red arrows, net decrease of >1000 reads in peak in carboplatin treated cells, respectively. **F,** RISC-bound sRNA Seed Tox graphs of PEO4 cells treated with 1 μg/ml carboplatin or water for 72 hrs. **G,** RISC-bound sRNA Seed Tox graphs of water treated PEO1 and PEO4 cells. Green arrows, net increase, and red arrows, decrease of >1000 reads in peak in PEO4 cells, respectively. **H,** RISC composition of PEO1 or PEO4 cells. Shown are piecharts of short-term (S-Carb) or long-term (RX) treated cells isolated from patients. Reads with a predicted 6mer seed viability of <20% are shown in red, >80% in green, and in between in grey. P values were calculated using pairwise comparisons of all reads in the two different treatments in two groups: reads with predicted viability >80% and <20% (blue stippled lines). In these areas, the most significantly changed miRNAs are labeled. The average miRNA content of RISC bound sRNAs across all A2780/PEO samples was 97.6%/97.8%, respectively. Length of green and red arrows corresponds to number of reads (inserts in A and E).

To determine whether the changes detected in cell lines treated *in vitro* can be found in the RISC of OC xenografts grown in mice and treated with Pt, we injected OVCAR5 tumors orthotopically into nude mice and treated them repeatedly with either PBS or carboplatin (Supplementary Fig. S6A). Tumors treated with carboplatin were in general smaller compared to those of PBS injected control animals (insert a in Supplementary Fig. S6B). Consistent with the cell line models of acquired Pt resistance, *in vivo* Pt treated Pt sensitive OVCAR5 tumors showed a reduction of toxic RISC-bound miRNAs with an increase in nontoxic miRNAs (Supplementary Fig. S6C). The most apparent increase of a nontoxic miRNA was in miR-194-5p, a miRNA recently reported to confer Pt resistance in OC (31) (Supplementary Fig. S6B, S6C). This increase correlated with the degree of Pt resistance of the tumors isolated from the carboplatin treated mice (insert b in Supplementary Fig. S6B) and supports our hypothesis that OC cells have different ways to acquire therapy resistance through upregulation of various miRNAs with nontoxic 6mer seeds.

### Uptake of Toxic siRNAs into the RISC of Platinum Tolerant Cells

To determine why some Pt resistant cells were more sensitive to toxic 6mer seed containing siRNAs, we transfected the A2780/A2780R cell pair with a low amount (1 nM) of the nontoxic siUAGUCG (siNT1) or either of the toxic siGGCAGU or siGGGGGC siRNAs. At this concentration, the latter two siRNAs were not toxic to the parental cells but significantly reduced cell growth of the Pt-R cells (Supplementary Fig. S7A). All transfected cells had large numbers of RISC-bound and total reads derived from the guide strand of the transfected siRNAs (Supplementary Fig. S7B). There were few reads derived from the blocked passenger strand (Supplementary Fig. S7C). We noticed an equal uptake of the nontoxic siRNA into Pt-S and Pt-R cells and their RISC, with a much higher uptake of the two toxic siRNAs in A2780R cells (Fig. 3A-D, Supplementary Fig. S7D-S7F).

**Figure 3.**
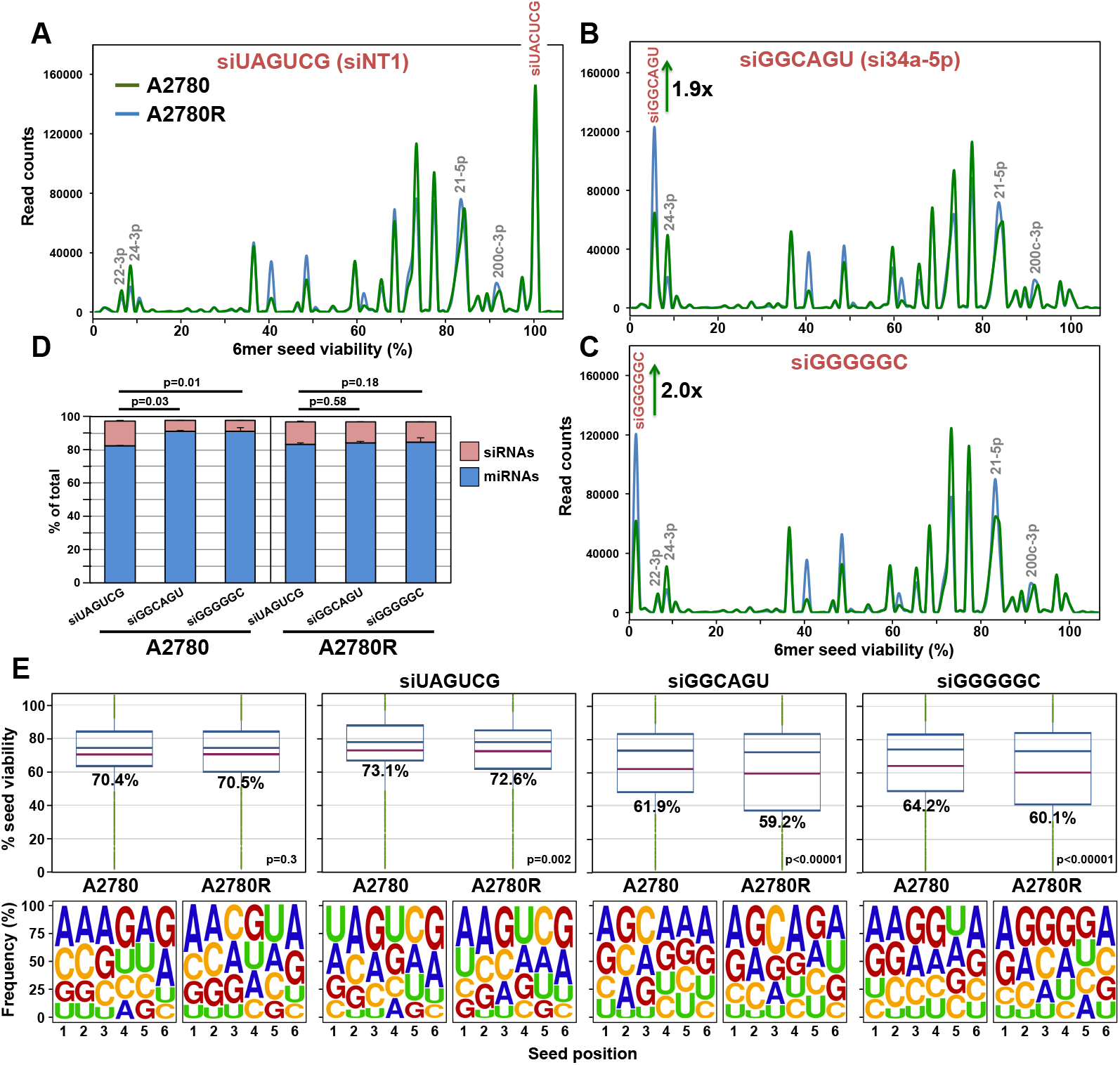
Platinum resistant, A2780R cells have more toxic miRNAs in their RISC than sensitive A2780 cells. **A-C,** Seed Tox graph of RISC bound miRNAs and exogenous siRNAs in A2780/A2780R cells 24 hrs after transfection with either the nontoxic siUAGUCG (siNT1), or the highly toxic siGGCAGU, or siGGGGGC. **D,** Percent miRNAs and exogenous siRNAs in the RISC of the siRNA transfected cells. Student’s test p-values are shown. **E,** Average Seed Tox (top) and seed composition (bottom) of all RISC-bound sRNAs in A2780/A2780R cells PBS treated or treated with the three siRNAs. P values at the top were calculated using a Wilcoxon rank test. The fraction of exogenous siRNAs and endogenous miRNAs was ~ 97% in all samples.

Based on these results, we were able to assess the ratio of toxic siRNA to RISC bound miRNA required to reduce cell growth. Because Ago k.o. cells are resistant to the effects of toxic siRNAs, we know that the toxicity is dependent on the presence of Ago2 in the RISC. The nontoxic siUAGUCG replaced about the same amount of miRNAs from the RISC in Pt-S and Pt-R cells, while the two toxic siRNAs were less efficient in replacing miRNAs in the Pt-S cells (Fig. 3D). The overall seed toxicity of all RISC bound reads in cells transfected with the two toxic siRNAs significantly dropped in A2780R cells, resulting in a relatively small change in overall 6mer seed composition of RISC bound sRNAs between the A2780 and A2780R cells (Fig. 3E). These data suggest that a moderate shift of seed toxicity of about 10% or less is sufficient to reduce cell growth and/or cause cell death in response to Pt.

### Changes in Tumors of OC patients Caused by Platinum Based Therapy

To determine whether changes in toxic and nontoxic 6mer seed containing miRNAs are associated with Pt resistance in OC patients, we studied a well characterized group of Pt sensitive and Pt resistant patients with FIGO stage III/IV high grade serous OC who underwent primary debulking surgery followed by adjuvant chemotherapy with carboplatin and paclitaxel (Supplementary Table S3). Fresh frozen primary OC tissue specimens were subjected to an Ago pull down/smRNA Seq analysis. Two cohorts of patients were compared: Group I: Pt resistant; Group II: Pt sensitive (Fig. 4A). Because the RISC composition could be affected by the age of the patients, both groups were age matched (Supplementary Fig. S8A). miRNA enrichment in the pulled down RISC was highly efficient. Across all 21 samples, between 97.4% and 98.1% of all Ago bound reads were miRNAs. When ranking all highly abundant reads derived from miRNAs (sum >10,000 across all samples) according to the fold change in the Pt-S patients, we found that tumors in Pt-S patients had an enrichment of miRNAs with more toxic seeds while Pt-R tumors were enriched in miRNAs with *non*toxic seeds (Fig. 4B). This correlation was also apparent when all ~38,000 reads were ranked according to the predicted 6mer Seed Tox of each read (Supplementary Fig. S8B). A shorter time to relapse correlated with higher amounts of nontoxic RISC-bound miRNAs.

**Figure 4.**
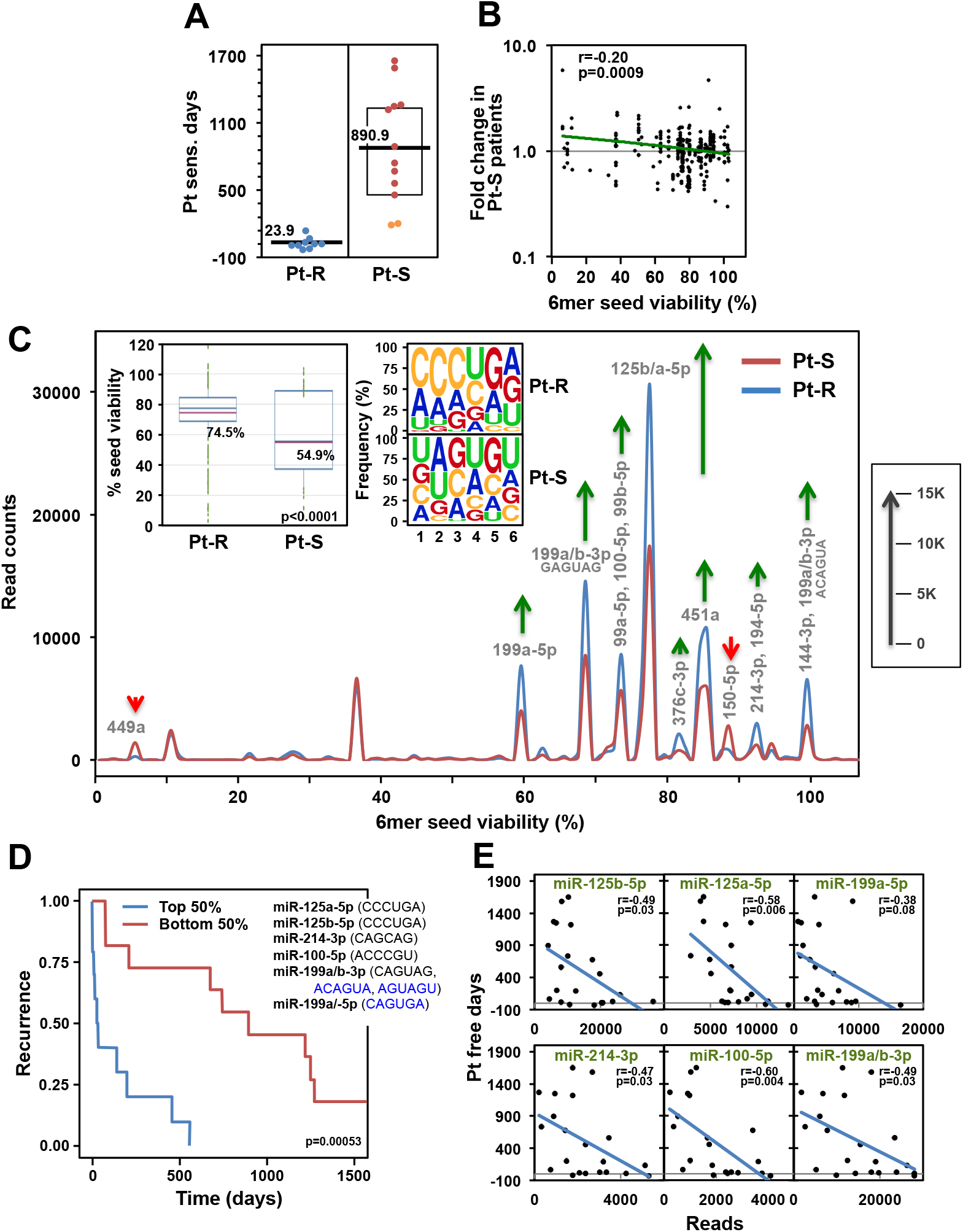
Lower Seed Tox of RISC-bound sRNAs in OC tumors is associated with increased resistance to chemotherapy. **A,** Pt sensitive days (days between the end of the last cycle of Pt-based chemotherapy and first relapse) in two groups of OC patients: Pt resistant (Pt-R) and Pt sensitive (Pt-S). Orange dots indicate two patients with intermediate (180 – 365 days since last treatment) Pt sensitivity. **B,** Pearson correlation between the fold change in Pt-S versus Pt-R patients and the 6mer Seed Tox (% viability in HeyA8 cells). Shown are miRNAs with a sum of >10,000 across all samples (representing 85.4% of all reads). miRNA content of this pull down was 97.4-98.1%. **C,** Seed Tox graph of RISC-bound miRNAs differentially expressed (p<0.05, >1.5x fold change) in Pt-S versus Pt-R patients. Green arrows, net increase, and red arrows, net decrease of >1000 reads in peak in Pt-R tumors, respectively. Inserts: *Left:* Average Seed Tox of all sRNAs enriched in either Pt-R or Pt-S patients. *Right:* matching seed compositions. **D,** Kaplan-Meier analysis of patients with the top and bottom highest RISC content of the average of the six miRNAs shown (including their 6mer seeds). Noncanonical seeds are in blue. **E,** Regression analysis of the six miRNAs in D plotted against the number of Pt sensitive days.

Comparing the significant differences in RISC-bound miRNAs between the two groups in a Seed Tox graph identified miR-449 and miR-195-5p as depleted and several miRNAs as enriched (e.g. miR-125a/b) in Pt-R tumors (Fig. 4C). That RISC-bound miRNAs enriched in the Pt-R tumors had a significantly higher seed viability than in the Pt-S tumors, was also reflected in the change in seed composition (insert in Fig. 4C). These data suggest that it is not an individual miRNA but a combination of nontoxic miRNAs that confers Pt resistance in human OC.

The average 6mer seed composition of total RISC-bound miRNAs in the tumors was different (Supplementary Fig. S9A). An unsupervised hierarchical cluster analysis of the reads with the ten most frequent 6mer seeds separated patient tumors into two subtypes (Supplementary Fig. S9B). One subtype contained seeds present in miR-200 family members, and the other subtype contained seeds found in miR-21-5p and members of the let-7-5p family. However, both subtypes were represented in Pt-S and Pt-R patients and therefore do not correlate with outcome.

When focusing on only the most highly expressed miRNAs in Pt-R patient tumors, we found six miRNAs to correlate with Pt resistance in OC patients (Supplementary Table S4). Grouping patients with the 50% highest and lowest combined reads of these six miRNAs separated Pt-S from Pt-R patients (Fig. 4D). These six highly abundant miRNAs also correlated individually with the Pt sensitive days (Fig. 4E). Individual Kaplan-Meier analyses with these six miRNAs showed that the separation is largely driven by four miRNAs (miR-125a-5p, miR-125b-5p, miR-100-5p, and miR-199-5p). Using the combination of these miRNAs allowed the best distinction between Pt-S and Pt-R patients (Supplementary Fig. S10).

### Analysis of Primary Tumors and Recurrences in Patients with Very Long Overall Survival

A comparison of matched primary tumors with recurrences is often not feasible in OC, because patients are usually only operated on once. However, we recently described a subgroup of patients who, for unknown reasons, maintain Pt sensitivity and/or have oligometastatic recurrences (32, 33). Because some of these patients undergo a secondary surgery, we were able to compare 6mer Seed Tox between the initial tumor and the recurrence (Fig. 5A and Supplementary Table S3). Two patients (#290 and #200) overlapped with the above analysis (Fig. 4); however, we analyzed a different tumor site, allowing us to test both the similarities between these tumors and the reproducibility of the Ago pull down/smRNA Seq analysis (Supplementary Fig. S11). The independent analyses of the two tumors from the same patients were more similar when compared to the average of the other primary tumors in the second set.

**Figure 5.**
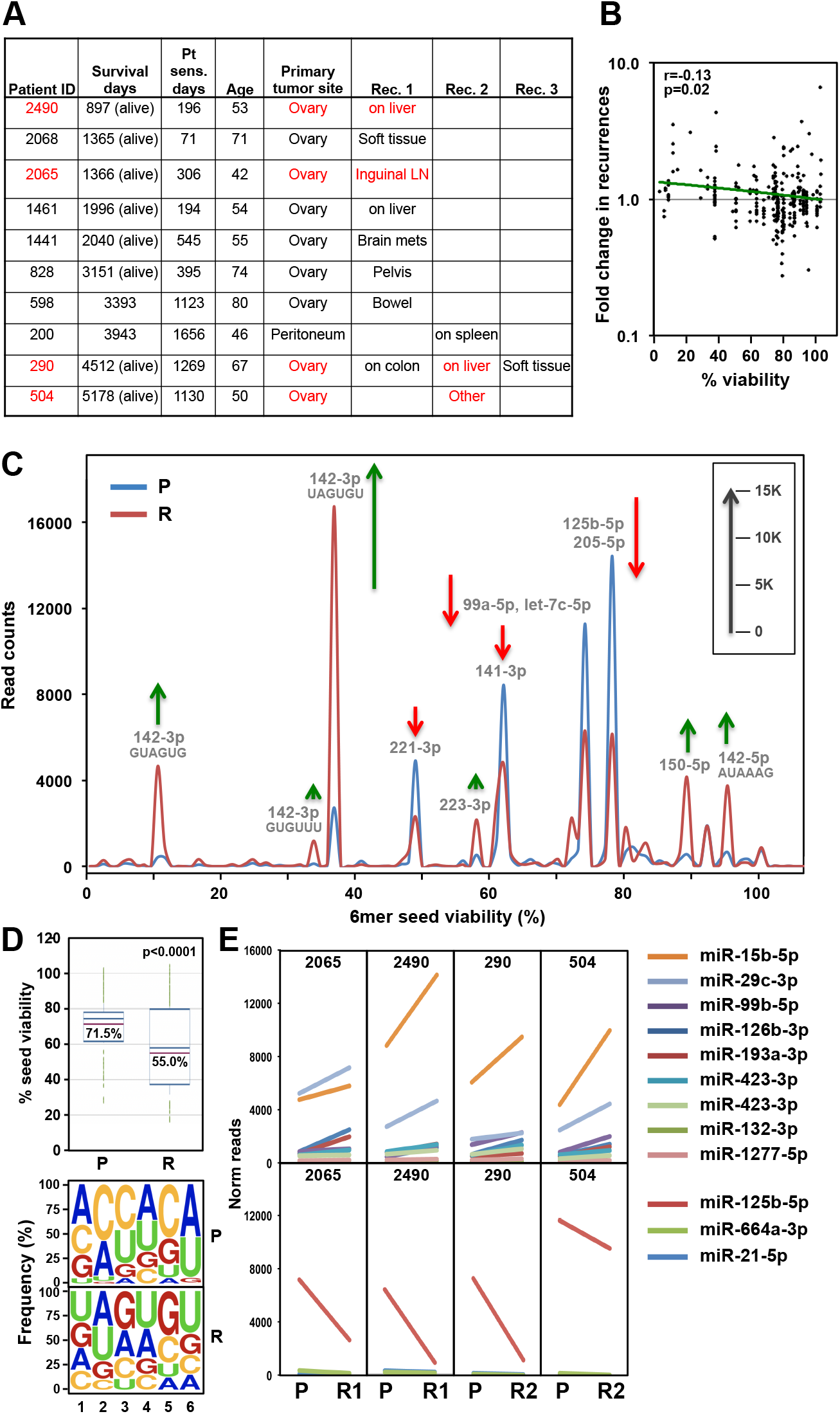
The RISC in recurrent OC tumors in a group of long-term survivors is enriched in miRNAs with toxic 6mer seeds compared to the primary cancer. **A,** List of 10 long-term survivor OC patients with sites of recurrences given. Patient pairs of a subgroup of four highly related primary and recurrent tumor pairs are shown in red (see Supplementary Fig. S12B). Survival days are as of 12/22/2020. **B,** Pearson correlation between the fold change in recurrences versus primary tumors and the 6mer Seed Tox (% viability in HeyA8 cells). Shown are miRNAs with a sum of >10000 across all samples (representing 81.4% of all reads). miRNA content of this pull down was 97.2-98.4%. **C,** Seed Tox graph of RISC-bound miRNAs differentially expressed (paired analysis, p<0.05, >1.5x fold change) in recurrences versus matched primary tumors in the subgroup of four patients. Note, miR-15b-5p was not detected in this analysis which used the EdgeR software package for to obtain paired p-values and it had a p-value of 0.057. Green arrows, net increase, and red arrows, net decrease of >1000 reads in peak in recurrent tumors, respectively. **D,** *Top:* Average Seed Tox of all sRNAs enriched in either primary tumor (P) or recurrences (R). *Bottom:* Matching seed compositions. **E,** Changes in RISC bound miRNA read numbers in recurrences versus primary tumors in the subgroup of patients. Shown are all miRNAs that are significantly up (top) or downregulated (bottom) (paired Student’s t-test p<0.05, >1.5x fold) in all four patients in the same direction.

Overall Seed Tox was higher in the recurrent tumors based on a comparison of the most abundant reads in the matched primary tumors (Fig. 5B). Integrating all reads of the primary tumors and the recurrences revealed that some miRNAs with toxic seeds were upregulated (e.g. miR-22-3p and the miR-15/16-5p family), while miRNAs with nontoxic seeds (miR-26a/b-5p, let-7-5p family, miR-143-5p, miR-125a/b-5p, miR-30-5p family) were downregulated in the recurrent tumors (Supplementary Fig. S12A). While the recurrences and primary tumors were from the same patients, we could not exclude the possibility that the difference in Seed Tox of RISC bound miRNAs was due to differences in the tumor microenvironment or the distinct anatomic locations of the 1^st^ and 2^nd^ surgery. An unsupervised hierarchical cluster analysis based on RISC-bound miRNAs identified the primary and recurrent tumors from four patients as highly related regardless of anatomic location, suggesting that the detected miRNAs were tumor-specific (Supplementary Fig. S12B). A Seed Tox graph was generated of the samples of these four tumor pairs (Fig. 5C). A few miRNAs with toxic seeds were upregulated in the recurrences, while several miRNAs (e.g. miR-125b-5p) with nontoxic seeds were downregulated. This resulted in a significant drop in average toxicity of all miRNAs enriched in recurrences (Fig. 5D). To identify the miRNAs driving these changes, we plotted all miRNAs that were significantly deregulated in the same direction in this subgroup of patients (Fig. 5E). In this analysis, in which we applied a paired Student’s T test in three out of 4 patients, the most abundant upregulated miRNA was miR-15b-5p, which can kill cancer cells through 6mer Seed Tox (22), and the most abundant downregulated nontoxic miRNA was, again, miR-125b-5p. Analysis of all patient tumor pairs suggests that the differences seen between primary tumors and matched recurrences in this rare patient population are due to intrinsic changes in the tumor cells. Tumors of “multiple Pt responders” (34), do not acquire Pt resistance through upregulation of nontoxic/protective miRNAs. Based on our data, we predict that the recurrences in these patients will be as Pt sensitive as the initial tumors.

## DISCUSSION

We have advanced a novel concept to explain how miRNA families target gene networks that regulate cell fate. This concept is not based on the activity of any individual miRNA or its targets, but on the conclusion that it is the incidence of 6mer Seed Tox in all miRNAs present that determines whether genes critical for cell survival are being silenced. We are now proposing that the balance between miRNAs with highly toxic seeds and miRNAs that carry nontoxic seeds determines treatment outcome in OC. Testing this concept required assessing position 2-7 of RISC bound miRNA guides. To obtain this information, we combined the method of Ago pull down/smRNA Seq with a specific bioinformatics pipeline to analyze 6mer seeds in short reads. Our method does not involve crosslinking of the miRNAs with their target mRNAs, since the primary goal of this study was not to identity miRNA targets, but to determine the seed content of the RISC in cancer cells and tumor tissue. Recent data suggest that many miRNAs detected in total RNA are part of a low-molecular-weight RISC that is not associated with either GW182 or mRNAs (30). By using a GW182 peptide to pull down Ago proteins, we may have preferentially isolated active RISC complexes, as only those active (high-molecular-weight) complexes are comprised of miRNA, Ago, mRNAs and GW182 (30). This is supported by our recent observation that the GW182 peptide could not efficiently pull down Ago in Drosha k.o. cells devoid of most miRNAs (21) despite unchanged Ago expression levels (18) It also provides an explanation for the large variation we found between total and RISC-bound miRNAs for a number of miRNAs. Ago proteins are stabilized by bound miRNAs and the number of mRNAs with targeted seed matches that bind to these miRNAs (29). Our finding that the two toxic siRNAs, siGGCAGU and siGGGGGC, are present in higher numbers and loaded into the RISC more efficiently than the nontoxic siUAGUCG when transfected into the A2780R cells, could be because the two toxic siRNAs target survival/housekeeping genes in this cell line (21).

We developed a way to visualize all significantly expressed miRNAs in one simple graph. The 6mer Seed Tox graph relies only on the predicted toxicity of every miRNA based on our screen of all 6mer seeds. miRNAs that are part of different families with entirely different biological functions may have similar seed toxicities and will be found in the same peaks. Based on these analyses, we are proposing that it is the balance between toxic and nontoxic 6mers embedded in miRNAs that determines whether cancer cells die or resist Pt-based chemotherapy (Fig. 6). We provide the first evidence that efficient Pt induced cell death is in part dependent on a functional RNAi system and that cells devoid of miRNAs (most of which contain nontoxic seeds) are highly sensitive to Pt induced cell death. The immediate reaction of OC cells to Pt is downregulation of miRNAs with nontoxic seeds and upregulation of miRNAs containing highly toxic 6mer seeds such as miR-22-3p and miR-24-3p (Fig. 6I). Both of these miRNAs are upregulated in human embryonic stem cells lines upon differentiation (35) and both have been shown to induce cell death when transfected into HCT116 cells (36). In contrast, OC cells that have acquired resistance after long-term Pt exposure show the opposite pattern: These Pt-R cells contain fewer toxic miRNAs and more nontoxic RISC-associated miRNAs (Fig. 6II). However, this resistance can be overcome by introducing artificial miRNAs with highly toxic seeds (e.g. siGGGGGC or siGGCAGU, Fig. 6III). Finally, when the nontoxic miRNAs are predominately one species (e.g. miR-21-5p in the A2780R cells) inhibiting this miRNA can re-sensitize cells to treatment (Fig. 6IV). Consistent with the *in vitro* data, tumors from patients with Pt resistance had more miRNAs with nontoxic seeds, while Pt sensitive patients had more miRNAs with toxic seeds. Confirming the proposed concept, recurrent tumors from patients that remain Pt sensitive had higher concentrations of toxic miRNAs in their RISC compared to the original primary tumor. In both analyses of patient tumors, the nontoxic miRNA most relevant to primary OC tumors was the abundant miRNA miR-125b-5p.

**Figure 6.**
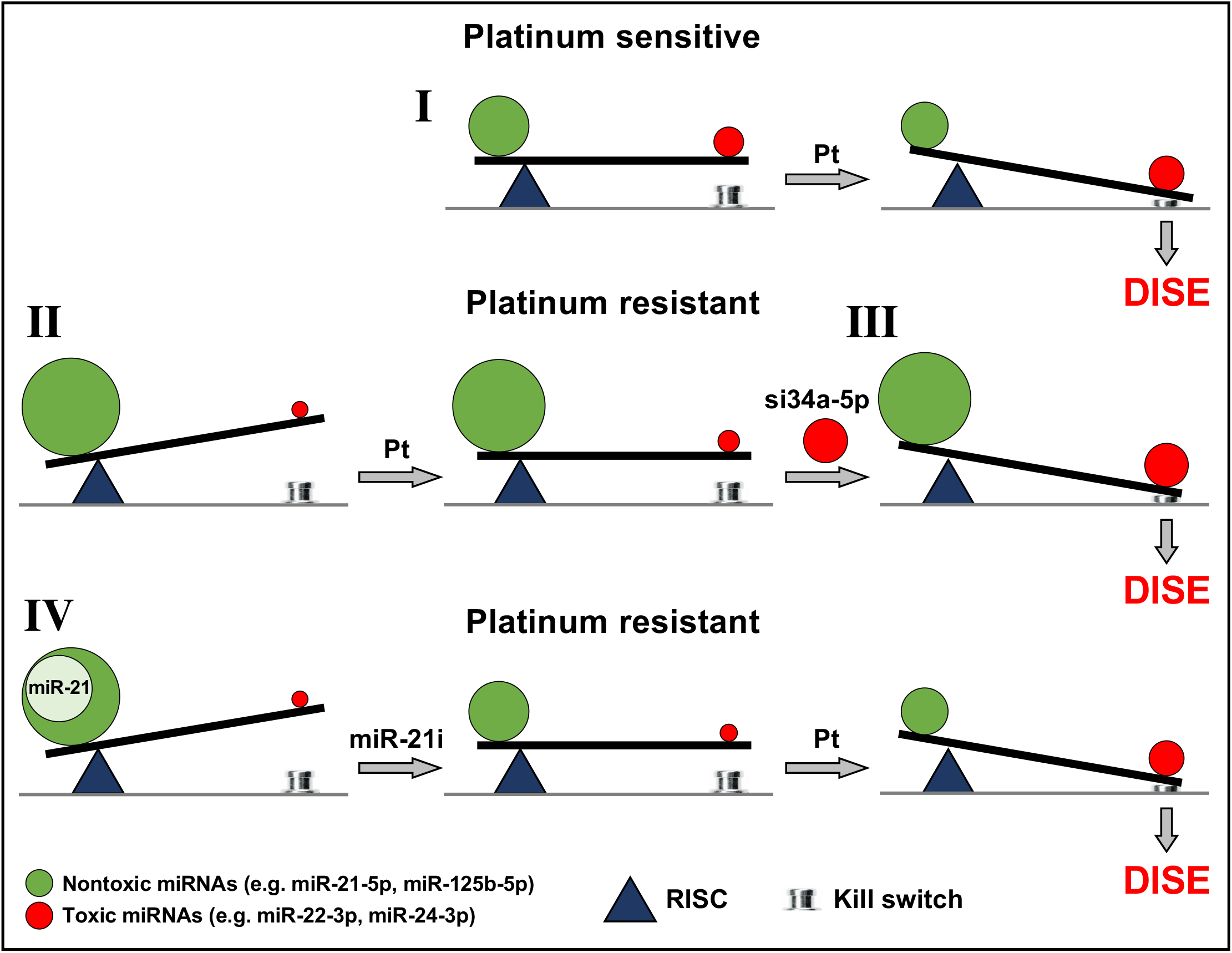
Scheme to illustrate the balance of toxic RISC bound miRNAs that determine platinum sensitivity. In constitutively Pt sensitive OC cells treatment with Pt results in a shift in the ratio of miRNAs with toxic and nontoxic seeds tipping the balance towards cell death (death induced by survival gene elimination, DISE) (I). In OC cells with acquired Pt resistance the shift toward toxic seed containing miRNAs is insufficient to trigger DISE (II). However, the protection by miRNAs with nontoxic seeds can be overcome by introducing exogenous miRNA mimetics (e.g. siGGCAGU = si34a-5p) that contain highly toxic seeds (III). Finally, inhibiting miRNAs with nontoxic seeds, as shown for overexpressed miR-21-5p (using a miR-21-5p inhibitor, miR-21i) in the A2780R cells, allows the Pt resistant cells (both constitutively as well as acquired) to regain sensitivity to Pt (IV).

While our study is based on a relatively small number of patients, this is the first analysis of primary human tumors based exclusively on RISC-bound miRNAs. Traditionally, analysis of miRNAs in human cancers is done using total RNA. It is based on the assumption that quantification of total miRNA amounts in cells or tissue allows the assessment of the functional relevance of expressed miRNAs. Yet, a recent study has shown that the level of Ago-bound miRNAs is a better indicator of miRNA activity than the total level of miRNA expression. The differential binding of miRNAs to the RISC was found to be cell type specific (29), and to be independent of GC content of miRNAs. However, it was strongly dependent on the abundance of complementary mRNA targets. Our data now suggest this is true of ovarian tumors. We identify a number of miRNAs that are preferentially associated with Ago proteins, some of which we then found to be associated with the response of OC cells to Pt-based chemotherapy.

Our data suggest that focusing on single miRNAs or even miRNA families will not result in a reliable marker for Pt resistant OC. We hypothesize that OC cells can acquire Pt resistance by upregulating any miRNA that carries a nontoxic 6mer seed. This hypothesis is consistent with the countless reports that provide evidence for the involvement of single miRNAs in Pt resistance in OC cell lines (31, 37–44) (Supplementary Table S1 and S2). Pt resistance in different OC cell line models and in OC patients may be driven by a variety of nontoxic miRNAs. This is supported by our smRNAseq data derived from different models (Table 1). We conclude that Pt resistance in each of the three models and in human patients may be driven by different nontoxic miRNAs reported to render OC cells Pt tolerant. Our cisplatin resistant A2780R utilize miR-21-5p, PEO4 cells are more tolerant to carboplatin due, in part, to upregulation of miR-130a-3p, and OVCAR5 cells exposed to prolonged treatment with carboplatin *in vivo* upregulate mostly miR-194-5p. Finally, in tumors from Pt-R patients, we found evidence for an upregulation of RISC-bound miR-214-3p and, most prominently, miR-125b-5p in the most Pt resistant patients. While any of these miRNAs could serve as targets for therapy and as biomarkers, it is entirely possible that different miRNAs will be identified which confer therapy resistance in other cell lines and patient analyses. What they may all share is the presence of a nontoxic seed. Artificial toxic 6mer seed containing miRNAs or siRNAs could therefore be used to eliminate OC and other cancers. Indeed, our data suggest that Pt resistant tumors may be at least as sensitive to these toxic siRNAs as chemotherapysensitive tumors.

**Table 1:** Different miRNAs are associated with Pt resistance in different models of OC and in OC patients. For each miRNA listed 6mer seed viability was determined in HeyA8 cells (6merdb.org). Only shown are miRNAs with average reads >1000 across all samples. Samples with average <1000 reads are labeled with /. nd, not detected = less than 10 reads across all samples. nd, not detected, less than 10 reads across all samples. In addition, miRNAs found to be enriched in Pt-R patients and to significantly correlate with Pt sensitive days are shown in the last column. Both Pearson coefficients and p-values are given. Only miRNAs are listed for which it has been demonstrated that overexpression of a miRNA mimic in Pt-S cells increases Pt resistance and inhibition of the same miRNA in isogenic Pt-R cells increases sensitivity. This was shown for miR-21 (37, 38), miR-125b (39), miR-130a and miR-374a (40), miR-149-5p (41), miR-214-3p (42), miR-216a-5p (43), miR-551b-3p (44), and miR-194-5p (31).

| miRNA | Seed viab. | A2780R vs S | PEO4 vs PEO1 | OVCAR5 + L-Carb | Corr. Pt-R vs Pt free days |
| --- | --- | --- | --- | --- | --- |
| 21-5p | 84.2% | **3.3** | 1 | 1.1 | 0.49 (p=0.026) |
| 125b-5p | 77.5% | 0.1 | 0.8 | 0.5 | **-0.49 (p=0.026)** |
| 130a-3p | 81.6% | 0.5 | **2.6** | 0.8 | -0.11 (ns) |
| 374a-5p | 88.1% | 1 | 1.1 | 1.2 | 0.07 (ns) |
| 149-5p | 63.9% | / | / | / | / |
| 214-3p | 78.1% | 1 | / | 1.1 | **-0.46 (p=0.034)** |
| 216a-5p | 64.9% | nd | nd | / | / |
| 493-5p | 75.6% | nd | nd | / | / |
| 551b-3p | 69.2% | / | / | / | / |
| 194-5p | 93.2% | / | / | **2.0** | / |

## METHODS

### Reagents and Antibodies

Carboplatin (#C2538) was purchased from Sigma-Aldrich, cisplatin (#1134357) was from Millipore Sigma, Lipofectamine RNAimax (#13778150) from Thermofisher Scientific. Primary antibodies for western blot: Anti-Argonaute-2 antibody (#ab32381) was purchased from Abcam; anti-Dicer (D38E7) antibody (#5362) from Cell Signaling; anti-β-actin antibody (#sc-47778) was purchased from Santa Cruz. The secondary antibody was a goat anti-rabbit IgG-HRP (#4030-05) from Southern Biotech.

### Cell Lines

Human OC cell lines, HeyA8, OVCAR5, were cultured in RPMI 1640 medium (Cell gro #10-040 CM), supplemented with 10% heat-inactivated fetal bovine serum (FBS) (Sigma-Aldrich # 14009C), 1% L-glutamine (Mediatech Inc), and 1% penicillin/streptomycin (Mediatech Inc); PEO1 and PEO4 cells were cultured in the above medium and supplemented with 2 mM sodium pyruvate (Mediatech Inc). 293T WT, 293T Dicer k.o. cells [Clone #2-20; (45)], HeLa control and HeLa Ago2 k.o. were a kind gift from Dr. Sarah Gallois-Montbrun (46), 293T Ago2 k.o. cells were provided by Dr. Klaas Mulder (47), and OC cell lines A2780 and A2780R (25) were all grown in DMEM medium (Gibco #12430054) supplemented with 10% FBS, 1% L-glutamine and 1% penicillin/streptomycin. All cell lines were authenticated by STR profiling and tested for mycoplasma using PlasmoTest (Invitrogen).

### Western Blot Analysis

Cells were lysed using Killer RIPA lysis buffer (150mM NaCl, 10 mM Tris HCl pH7.2, 1% sodium dodecyl sulfate (SDS), 1% TritonX-100, 1% Deoxycholate, 5 mM EDTA) and freshly added PMSF (1 mM) and Protease Inhibitor Cocktail (Roche #11836170001). Equal amounts of protein (30 μg) were resolved on a 10% SDS-PAGE gel and Western blot was performed as previously described (18). Visualization of protein bands was performed using the SuperSignal™ West Dura Extended Duration Substrate (Thermo Fisher Scientific #34076). Antibodies diluted in 5% dry milk powder in 0.1% Tween-20/TBS (TBST): anti-Argonaute-2 (1:1000), anti-β-actin (1:5000), anti-rabbit IgG HRP (1:5000). Anti-Dicer antibody was diluted (1:1000) in 5% BSA in TBST.

### Assessing Cell Growth and Viability Assay

For transfection in 96-well plates to monitor cell growth with IncuCyte Zoom (Essen Bioscience), 50 μl Optimem (Fisher Scientific #331985088) mix containing RNAimax and 10 nM siRNA (siNT1, siGGCAGU or siGGGGGC) was used. RNAimax amounts per 96-well were optimized for each cell line used: A2780 (0.2 μl), PEO1/4 (0.3 μl), 293T (0.1 or 0.2 μl), OV81.2 (0.2 μl). Seeding densities for reverse transfection with siRNAs or Pt treatment in 96-well plates were also optimized for each cell line used: A2780/A2780R and OV81.2/OV81 CP40 (1500 cells), PEO1/PEO4 and 293T (1500 cells). The following siRNA oligonucleotides were obtained from Integrated DNA Technology and annealed per the manufacturer’s instructions: siUAGUCG (siNT1) sense: mUmGrGrUrUrUrArCrArUrGrUrCrGrArCrUrArATT siUAGUCG (siNT1) antisense: rUrUrArGrUrCrGrArCrArUrGrUrArArArCrCrAAA siGGCAGU (si34a-5p^Seed^) sense: mUmGrGrUrUrUrArCrArUrGrUrArCrUrGrCrCrATT siGGCAGU (si34a-5p^Seed^) antisense: rUrGrGrCrArGrUrArCrArUrGrUrArArArCrCrAAA siGGGGGC sense: mUmGrGrUrUrUrArCrArUrGrUrGrCrCrCrCrCrATT siGGGGGC antisense: rUrGrGrGrGrGrCrArCrArUrGrUrArArArCrCrAAA

For treatment with cisplatin or carboplatin, cells were seeded in triplicates in 96-well plates and treated with the indicated drug concentrations. Cell growth was monitored using IncuCyte Zoom live-cell imaging system (Essen Bioscience) with a 10x objective. The confluency curves were generated using the IncuCyte Zoom software (version 2015A). A viability assay that measures the level of ATP within cells was done in 96-well plates. Briefly, 72 hrs post-treatment with drugs or 96 hrs post reverse transfection with siRNAs, media in each well was replaced with 70 μl fresh medium 70 μl of Cell Titer-Glo reagent (Promega #G7570) was added. The plates were covered with aluminum foil and shaken for 5min and then incubated for 10 min at room temperature before the luminescence was read on a BioTek Cytation 5. IC50 values for cisplatin or carboplatin in a viability assay comparing Pt sensitive with Pt tolerant cells were determined using GraphPad Prism 6 software by logarithm normalized sigmoidal dose curve fitting.

### miR-21-5p Inhibition and Real-time PCR

Real-time PCR was performed for miR-21-5p in A2780R cells infected with lentivirus from Zip control (pLenti-III-miR-GFP control vector from Applied Biological Materials #m001) or Zip-21 (LentimiRa-GFP-has-miR-21-5p vector, Applied Biological Materials #mh10276). Briefly, 25 ng total RNA was used to make cDNA using the High-Capacity cDNA Reverse Transcription Kit (Applied Biosystems #4368814). The qPCR was then done using the Taqman Gene Expression Master Mix (ThermoFisher Scientific #4369016) as previously described (48) using the following primers: hsa-miR-21 (Thermofisher #0003970) and Z30 (Thermofisher #001092). The relative expression of miR-21-5p was normalized to the level of Z30. Statistical analysis was performed using Student’s T-test.

### In Vivo Treatment of OVCAR5 Cells with Carboplatin

Female (6-8 weeks old) athymic nude mice (*Foxn1^nu^*, Envigo) were injected intraperitoneally (i.p.) with 2 million OVCAR5 OC cells to induce tumors. After 2 weeks inoculation (49), mice were treated i.p. with PBS (control, mice ID: 791, 792, 793, 794, n=4), or 25 mg/kg carboplatin (n = 5) in the following treatment regimens: once-a-week for 3 weeks (stage 1, n=2, mice ID: 795, 797); 3-week carboplatin followed by two-week no drug recovery (Stage 2, n = 1, mice ID: 796); additional two weeks (5 weeks carboplatin in total) carboplatin after recovery (Stage 3, n = 2, mice ID: = 798, 800). Xenograft tumors were collected 1 week after carboplatin treatment and processed for cancer cell isolation and RNA extraction. Tumors were mechanically and enzymatically dissociated in Dulbecco’s modified Eagle’s medium/F12 (Thermo Fisher Scientific, Ref# 11320) containing collagenase (300 IU/ml, Sigma-Aldrich, Cat# C7657) and hyaluronidase (300 IU/ml, Sigma-Aldrich, Cat# H3506) for 2 hours at 37°C. Red blood cell lysis used RBC lysis buffer (BioLegend, Cat# 420301), followed by DNase (1 mg/ml, Sigma Aldrich, Cat# DN25) treatment and filtering through a 40 μm cell strainer (Fisher Scientific, Cat#NC0147038) to produce single cell suspension. EasySep™ Human Epcam positive selection kit (Stemcell Technologies, Cat# 17846) was used on the single cell suspension for Epcam positive OC epithelial cell selection following manufacturer instructions.

### Small Total RNA Seq

For small RNA-seq experiments, cells were lysed using Qiazol. For all RNA-Seq samples, a DNAse digestion step was included using the RNAse-free DNAse set (Qiagen #79254). Total RNA was isolated using the miRNeasy Mini Kit (Qiagen # 74004). The quality of RNA was determined using an Agilent Bioanalyzer. The RNA library preparation and subsequent sequencing on Illumina HiSEQ4000 was then done by the NU-Seq Core at Northwestern University.

Raw read sequences were trimmed with trim_galore to remove all standard Illumina adapters, leaving reads at least 6 bp in length. In addition, any reads containing the substring GTCCGACGATC followed by 3 to 5 nucleotides were discarded. This removed any remaining 5’ adapter sequences and ensured that only RISC bound sRNAs were analyzed, which was crucial for determining position 2-7 of each read. Trimmed reads were then sorted, uniq’d, counted. A six nucleotide Unique Molecular Identifier (UMI) was removed from each read, and the numbers of reads associated with the same core read sequence and any UMI were summed to generate the raw read count for a particular sample. Counts were tabulated for each read and sample and normalized per million reads in the column (sample) sum. Reads were blasted (blastn-short) against custom blast databases created from all processed human miRNAs (miRbase 22.1) and from the most recent RNA world database (human_and_virus_vbrc_all_cali_annoDB_sep2015) obtained from the Tuschl lab at Rockefeller University. Processed miRNA hits were filtered for 100% identity, and a match length of at least 18 bases. RNAworld hits were filtered for at least 95% identity, with the match starting within the first 9 bp of the sequencing reads. These filtered blast results were used to annotate reads with their likely source, and when no blast hit passed these filters, the read was annotated “noMatch”. Reads were also associated with their 6mer seed (positions 2-7) and with the previously identified seed toxicity determined in human OC cells HeyA8 (21) (6merdb.org). For tractability, only reads with a total count of at least 6 CPM summed across all samples were retained for further analyses. To identify reads enriched in a particular group of samples, we used the R package EdgeR with the un-normalized read counts, annotating the resulting reads with seed toxicity and blast-based annotation as described above.

### Ago Pull-Down and Subsequent Small RNA-Seq

For Ago pull-down experiments, transfections or treatments described above for 96-well plates were scaled up for 150 mm dishes. At the required time point for pull-down, cells were washed with PBS and 10 million cell pellets were frozen at −80 °C until ready to be lysed using 1 ml NP40 lysis buffer [50 mM Tris pH 7.5, 150 mM NaCl, 5 mM EDTA, 0.5% NP-40, 10% (v/v) glycerol, 1 mM NaF; supplemented with 1:200 EDTA-free protease inhibitors (Millipore #539134) and 1:1000 RNaisin Plus (Promega #N2615) before use]. Mouse or patient tumor lysates were prepared by first chopping 250 mg tissue with a clean razor blade and then using a Dounce homogenizer containing 1 ml NP40 lysis buffer. In patient tumors for which 250 mg tissue was not available, at least 100 mg tissue was used for pull down. The tissue was homogenized by passing the pestle up and down the cylinder several times while keeping the homogenizer cool on ice. Cell or tissue lysates were then incubated on ice for 15 min, vortexed, and then centrifuged at 20,000 g for 20 min. The lysates were then transferred to siliconized microcentrifuge tubes (low-binding, Eppendorf #022431021), and Ago1-4 were pulled down using 500 μg of Flag-GST-T6B peptide (50) and with 80 μl anti-Flag M2 Magnetic beads (Sigma #M8823) for 3 hrs on a rotor at 4°C. The precipitate was washed three times in NP40 lysis buffer and during the last wash, 10% of the beads were removed and incubated at 95°C for 5 min in 2x SDS-PAGE sample buffer. The efficiency of the pull down was determined by running these samples on a 10% SDS-PAGE gel, transferring them to nitrocellulose membrane and immunoblotting against Ago2 (Abcam #32381). 500 μl TRIzol reagent was added to the remaining beads and RNA was extracted using the manufacturer’s instructions. The RNA pellet was dissolved in 20 μl water. Half of the RNA sample was dephosphorylated with 0.5 U/μl of CIP alkaline phosphatase at 37°C for 15 min and then radiolabeled with 0.5 μCi γ-^32^P-ATP and 1 U/μl T4 PNK kinase for 20 min at 37°C. The RNAs interacting with Ago1-4 were visualized on a 15% Urea-PAGE. The other half of the RNA sample was used for a small RNA library preparation, as previously described (51). Concisely, RNA was ligated with 3’-adenylated adapters and separated on a 15% denaturing Urea-PAGE. The RNA corresponding to insert size of 19-35 nt was eluted from the gel, ethanol precipitated and then ligated with the 5’ adapter. The RNA samples were then separated on a 12% Urea-PAGE, extracted from the gel and reverse transcribed using Superscript III reverse transcriptase (Invitrogen #18080-044) and the cDNA was amplified by PCR. The cDNA was sequenced on Illumina Hi-Seq 4000. 19nt RNA size marker: rCrGrUrArCrGrCrGrGrGrUrUrUrArArArCrGrA 35nt RNA size marker: rCrUrCrArUrCrUrUrGrGrUrCrGrUrArCrGrCrGrGrArArUrArGrUrUrUrArArArCrUrGrU The following set of eight 3’ adenylated adapters was used, each containing a unique 6mer barcode sequence at the 5’ end (underlined). These barcodes were used to separate the reads into individual pull-down samples before proceeding with the small RNA pipeline as described above. For ease of use, the sequence files uploaded to GEO have already been split into reads from individual experiments:

Adapter 1-rAppNN<u>CTGACA</u>TGGAATTCTCGGGTGCCAAGG-L

Adapter 2-rAppNN<u>ACTAGC</u>TGGAATTCTCGGGTGCCAAGG-L

Adapter 3-rAppNN<u>GTACGT</u>TGGAATTCTCGGGTGCCAAGG-L

Adapter 4-rAppNN<u>TGTACG</u>TGGAATTCTCGGGTGCCAAGG-L

Adapter 5-rAppNN<u>CAGCAT</u>TGGAATTCTCGGGTGCCAAGG-L

Adapter 6-rAppNN<u>TCATAG</u>TGGAATTCTCGGGTGCCAAGG-L

Adapter 7-rAppNN<u>ATAGTA</u>TGGAATTCTCGGGTGCCAAGG-L

Adapter 8-rAppNN<u>GATGCT</u>TGGAATTCTCGGGTGCCAAGG-L

5’ adapter with unique molecular identifiers (UMI): GTTCAGAGTTCTACAGTCCGrArCrGrArUrCrNrNrNrN 5’ adapter without UMI: rGrUrUrCrArGrArGrUrUrCrUrArCrArGrUrCrCrGrArCrGrArUrC (used only in the experiment shown in Fig. 2).

RT primer sequence: GCCTTGGCACCCGAGAATTCCA

Four 3’ PCR primers were used each containing a unique index (underlined) recognized by Illumina:

Primer1:CAAGCAGAAGACGGCATACGAGAT<u>CGTGAT</u>GTGACTGGAGTTCCTTGGCACCCGAGAATTCCA

Primer2:CAAGCAGAAGACGGCATACGAGAT<u>ACATCG</u>GTGACTGGAGTTCCTTGGCACCCGAGAATTCCA

Primer3:CAAGCAGAAGACGGCATACGAGAT<u>GCCTAA</u>GTGACTGGAGTTCCTTGGCACCCGAGAATTCCA

Primer4:CAAGCAGAAGACGGCATACGAGAT<u>TGGTCA</u>GTGACTGGAGTTCCTTGGCACCCGAGAATTCCA

5’PCR primer:

AATGATACGGCGACCACCGAGATCTACACGTTCAGAGTTCTACAGTCCGA

### The Seed Tox graph

Reads were aggregated (read numbers collapsed) first by the seed sequence (position 2-7 of each read) and then by the miRNA and RNA World RNA that the read aligned to (Blast search see above). Each seed sequence had an assigned percent cell viability from the 4096 siRNA screen in the OC cell line HeyA8 ((21) and 6merdb.org), the aggregated read numbers were further binned in RStudio based on the HeyA8 cell viability in 1% increments. Seed tox graphs (number of reads on Y axis and 6mer seed viability on X axis) were then generated using Excel with smoothing enabled. The file with the collapsed data was used to identify the most abundant miRNAs that make up each peak in the 6mer Seed Tox graph, When a peak consisted of more than one abundant miRNA (>5000 or >1000 reads) miRNAs were listed in the order of abundance.

### Analysis of smRNAs in Normal Tissues

Data of total smRNA were retrieved (accession #GSE11879) and analyzed from 9 human tissues. To compare different tissues and consistently identify and label the most abundant miRNAs, all 9 data sets were normalized to 1,000,000 reads. For each tissue, miRNAs were labeled in Seed Tox graphs with >5000 norm average reads. When a peak contained more than one miRNA they are shown in the order from highest to lowest abundance.

### Seed and Tox Analysis

To determine the average Seed Tox of all reads in a data set or to determine the average 6mer seed composition of reads in a data set, numbers for all individual reads were divided by a factor that reduced the number of the most abundant read to less than 1000. In a comparison analysis, significantly up- or downregulated reads were processed separately. Associated toxicity and 6mer seed sequences for each read were then multiplied by this factor using a Perl script. The resulting read numbers were plotted as a box plot using the chart function of StatPlus:Mmac Pro v.7.3.3. using default settings and p values of deregulated seed toxicities were determined using a Wilcoxon rank test. Expanded seed sequences were plotted using Weblogo (http://weblogo.berkeley.edu) as a custom frequency plot.

### Comparison of Total and RISC Bound sRNAs

RNA Seq data sets were used for either total smRNA (DISE26 for A2780 and GSM3029765/GSM3029766 for HeyA8) or Ago bound smRNAs (DISE18 for A2780 and GSM3029217/GSM3029218 for HeyA8). To compare the amount of total versus RISC bound miRNAs all four data sets (with two duplicates each) were combined into one file using R. Each read column was renormalized to 1,000,000 reads. All reads were isolated with a length of 19-25 nt before generating Seed Tox graphs.

### Patient Samples and Analysis

All patient samples (described in Supplementary Table S3) were collected at the University of Chicago Medical Center following approval from the Institutional Review Board, and after obtaining written informed consent from the patients. All patients included in the study underwent primary debulking surgery for advanced stage metastatic high grade serous OC by a gynecologic oncologist. No patients who underwent neo-adjuvant treatment were included, The tissue was snap frozen at the time of surgery and stored at −80C until further use. Clinico-pathologic data was collected prospectively and updated every three months based on clinical exam, CA-125, or imaging.

### Analysis of Correlation Between Patient Groups and Seed Tox

Two cohorts of patients were compared: Group I: 9 patients were Pt refractory or resistant (recurrence <180 days after the end of chemo = Pt-R patients), Group II was platinum sensitive, experiencing a late recurrence (>180 days) or no recurrence (= Pt-S patients). Total norm count data were used. Reads were aggregated according to reads and blasted miRNAs. Patients were either separated into the two groups (Pt-R and Pt-S) compared or read associated toxicity was correlated using Pearson correlation (including p values) with Pt sensitive days as given in Supplementary Table S3.

### Identification of The Most Abundant miRNAs Enriched in Pt-R Patients

Total norm count data after adapter removal were used. Reads were aggregated according to reads and blasted miRNAs. All reads with an average read number of 1000 or more across all samples were used for this analysis. Student’s t-test (p<0.05) was used to identify all miRNAs that were enriched in the Pt-R patients (>1.5x).

### Unsupervised Hierarchical Cluster Analysis and Heat Maps

Either the collapsed reads of the ten most abundant miRNAs (carrying the same 6mer seed) (Supplementary Fig. S9B) or the most abundant miRNAs (>10,000 Ave reads across all samples) (Supplementary Fig. S12B) were hierarchically clustered and plotted in R according to pairwise Pearson correlation.

### Kaplan-Meier Analysis

Samples were divided into two groups according to expression of particular seeds/miRNAs, and survfit/ggsurvplot was used in R to create Kaplan-Meier curves contrasting the time to recurrence between the two groups.

### Statistical Analyses

IncuCyte experiments were performed in triplicate, and the data expressed as mean ±standard error. Continuous data were compared using t-tests for two independent groups. For evaluating continuous outcomes over time, two-way ANOVA was performed using the Stata1C software. Treatment condition was used as a component of primary interest and time as a categorical variable. When controls were very different between different treated cells, a binomial test was used for comparisons of single proportions to hypothesized null values (21).

## Data Availability

smRNA Seq data were deposited at GEO under the GSE series GSE165148. Data can be accessed using reviewer access token ojsdemaabzytdgx.

## Author Contributions

**M. Patel:** Performed experiments, wrote the manuscript. **Y. Wang**: Performed experiments. **A.E. Murmann:** Performed experiments. **E.T. Bartom:** Performed bioinformatics analyses. **R. Dhir:** Provided OC patient material. **K. Nephew:** Contributed a cell line and expertise. **D. Matei:** Provided cell lines, expertise and supervised the animal experiments. **E. Lengyel:** Provided OC patient material, wrote the manuscript. **M.E. Peter:** Conceived the project, supervised research, and wrote the manuscript.

## Acknowledgments

We are grateful to Drs. Gallois-Montbrun and Mulder for providing Ago2 k.o. cell lines.

