## Supplementary Figures and Tables for "The balance between toxic versus nontoxic microRNAs determines platinum sensitivity in ovarian cancer"

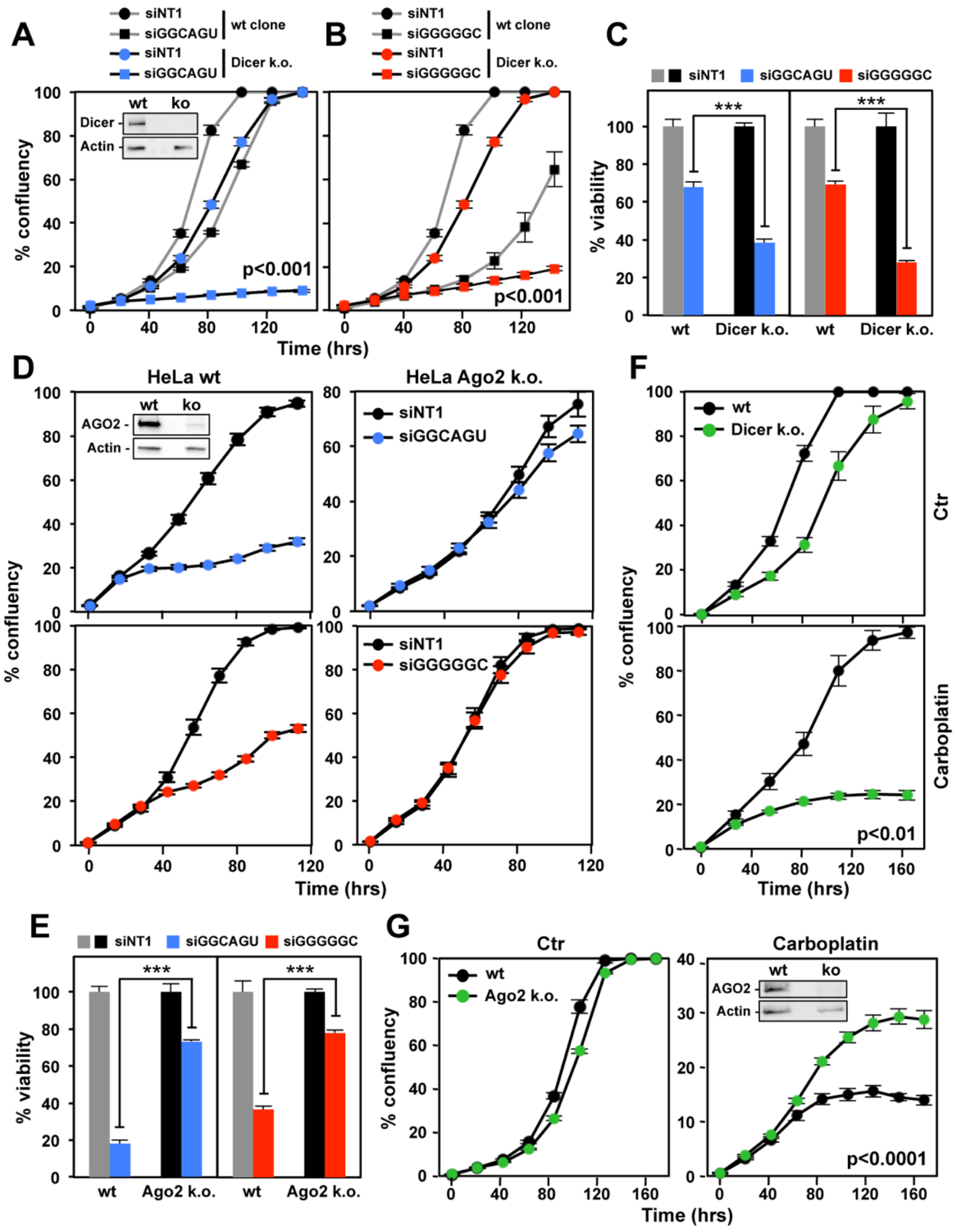


**Supplemental Figure S1.** Cell death induced by both toxic 6mer containing siRNAs and carboplatin involves RNAi. **A** and **B,** Confluency over time of 293T cells (wt or Dicer k.o.) transfected with 10 nM of siNT1, siGGCAGU or siGGGGGC. Insert: Western blot of Dicer in wt and k.o. cells. **C,** Viability of the cells in A and B 96 hrs after transfection with 10 nM of the siRNAs. **D,** Confluency over time of HeLa cells (wt or Ago2 k.o.) transfected with 10 nM of siNT1, siGGCAGU or siGGGGGC. Insert: Western blot of Ago2 in wt and k.o. cells. **E,** Viability of the 293T (wt or Ago2 k.o.) cells 96 hrs after transfection with 10 nM of the siRNAs. **F,** Confluency over time of 293T (wt or Dicer k.o.) treated with 3 μg/ml carboplatin. **G,** Confluency over time of 293T (wt or Ago2 k.o.) treated with 10 μg/ml carboplatin. Insert: Western blot of Ago2 in wt and k.o. cells. Each data point (A, B, D, F & G) represents mean ± SE of at least three replicates. Each bar (C, E) represents ± SD of three replicates. P-values were calculated using binomial distribution tests (A, B, F), ANOVA (G), or student's T-test (C, E). *** p<0.0001.

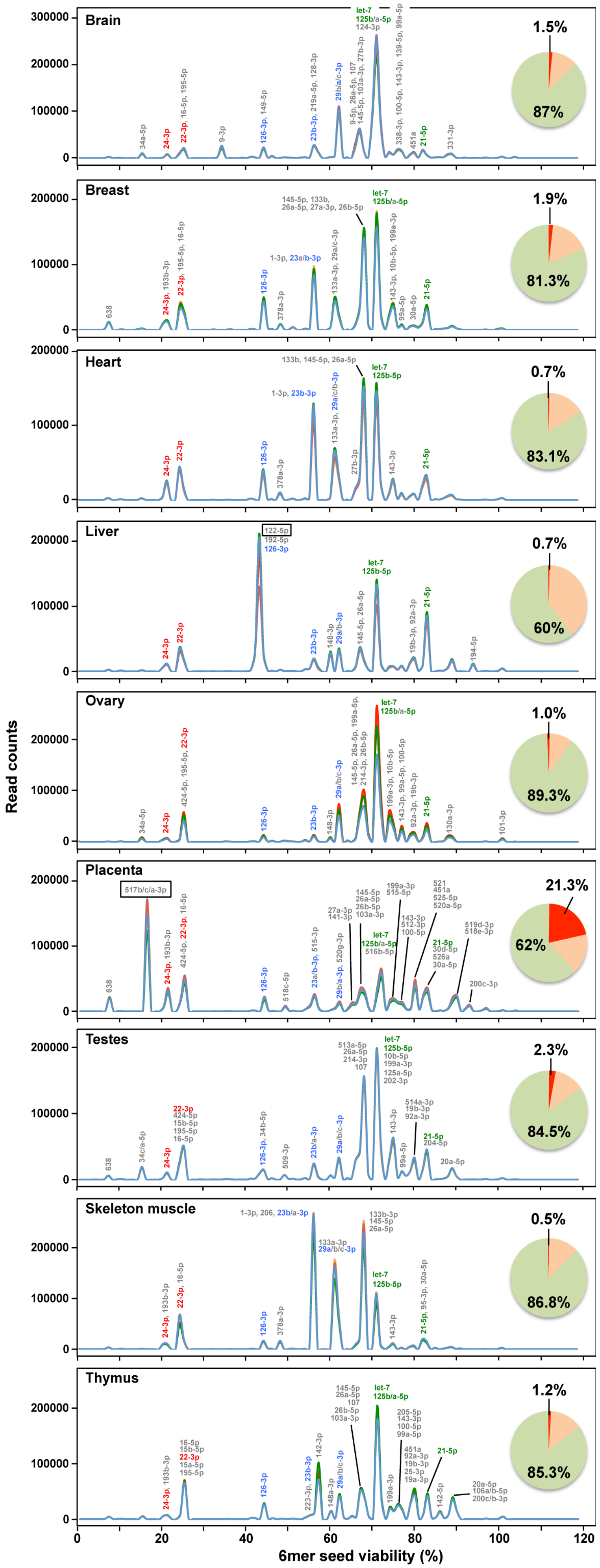


**Supplemental Figure S2.** Most tissues express predominantly nontoxic miRNAs**.** miRNA Seed Tox plots across 9 different tissues. In each case the cumulative read numbers are on the Y axis and the predicted toxicity on the X axis. Toxicity (shown as % viability) was the average of three human cell lines determined in a screen of all 4096 6mers in a neutral siRNA backbone (1). miRNAs significantly expressed in all tissues are labeled in color (red = highly toxic, blue = intermediately toxic, and green = nontoxic). When a peak is labeled with multiple miRNAs, the most abundant one is listed first. Pie chart insert: Abundance of miRNAs with seeds of the following predicted viabilities: red: <20%; corral: >=20 <50%; green: >50%. Data were normalized to 1 million reads and all miRNAs were labeled that have more than 5000 normalized average reads across five (for brain, breast, heart, liver, ovary, placenta, and skeleton muscle) or four (for testes and thymus) individual samples, respectively. Some established tissue specific miRNAs are boxed. GSE11879 was the source of the smRNA Seq data analyzed.

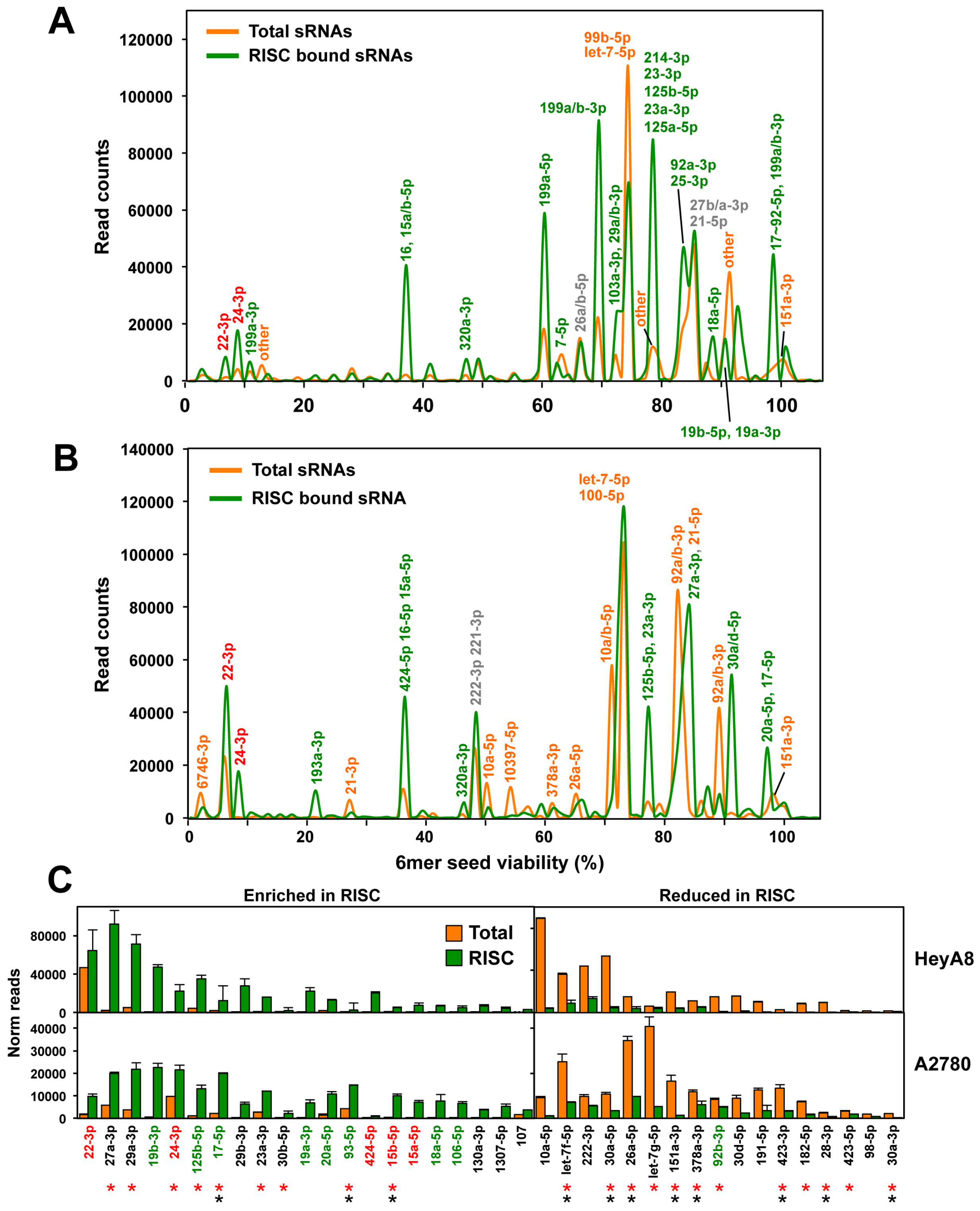


**Supplemental Figure S3.** Selective uptake of miRNAs into the RISC of OC cells. **A** and **B,** miRNA Seed Tox plots of either RISC-bound or total miRNAs in A2780 (A) and HeyA8 (B) cells. miRNAs that contribute to peaks with >5000 reads are labeled. For each labeled peak miRNAs are listed in the order of abundance. The percentage of miRNAs bound to the RISC was higher than the percentage of miRNAs in the total short (s)RNA population (79%/88.2% total and 97.6%/96.3% RISC bound in A2780 and HeyA8 cells, respectively). The seed viability (X axis) for HeyA8 cells was used (6merdb.org). **C,** All miRNAs with an average read number of >1000 in either cell line and >1.5 fold enriched (left) or reduced (right) in the RISC ranked according to the combined read numbers (total and RISC). miRNAs with toxic seeds are shown in red and putative protective miRNAs with nontoxic seeds are shown in green. miRNAs that have been reported before to be enriched or depleted in the same way in either 293T (red asterisks) or A549 (black asterisks) (2) are labeled.

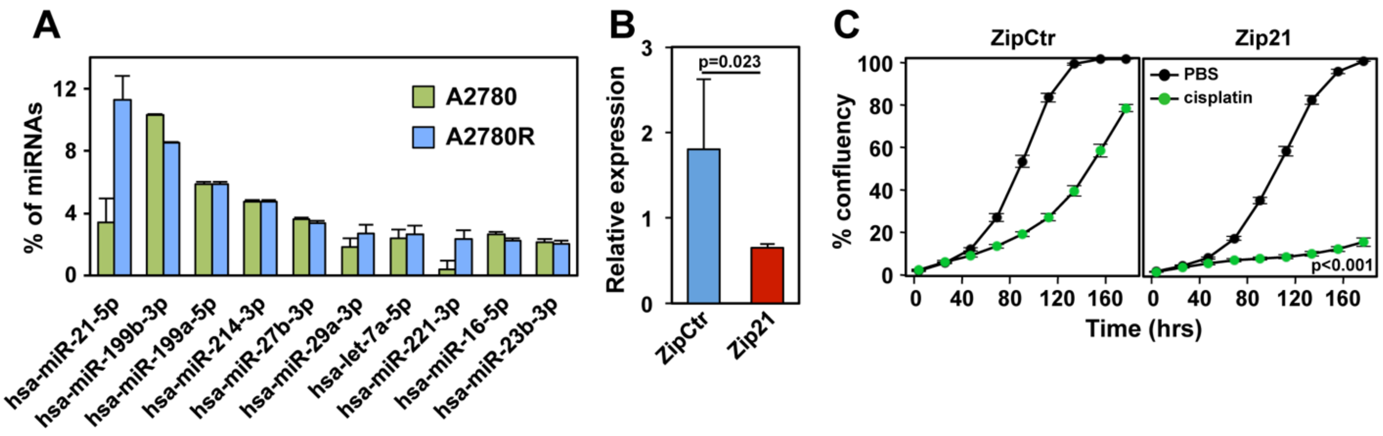


**Supplemental Figure S4.** miR-21-5p protects A2780R cells from platinum toxicity. **A,** Relative abundance (% of total reads) of the ten most highly expressed miRNAs in A2780 and A2780R cells (ranked according to highest expression in A2780R cells). Shown is the variance between duplicates. **B,** Real time PCR quantification of miR-21-5p in A2780R cells infected with control Zip vector or the miR-21 inhibitory lentivector Zip21. P-value from Student's T-test is shown. **C,** Confluency over time of ZipCtr or Zip21 expressing cells treated with either PBS or 5 μM cisplatin. P-value was calculated using a binomial distribution test. Experiments in B and C were done in triplicates.

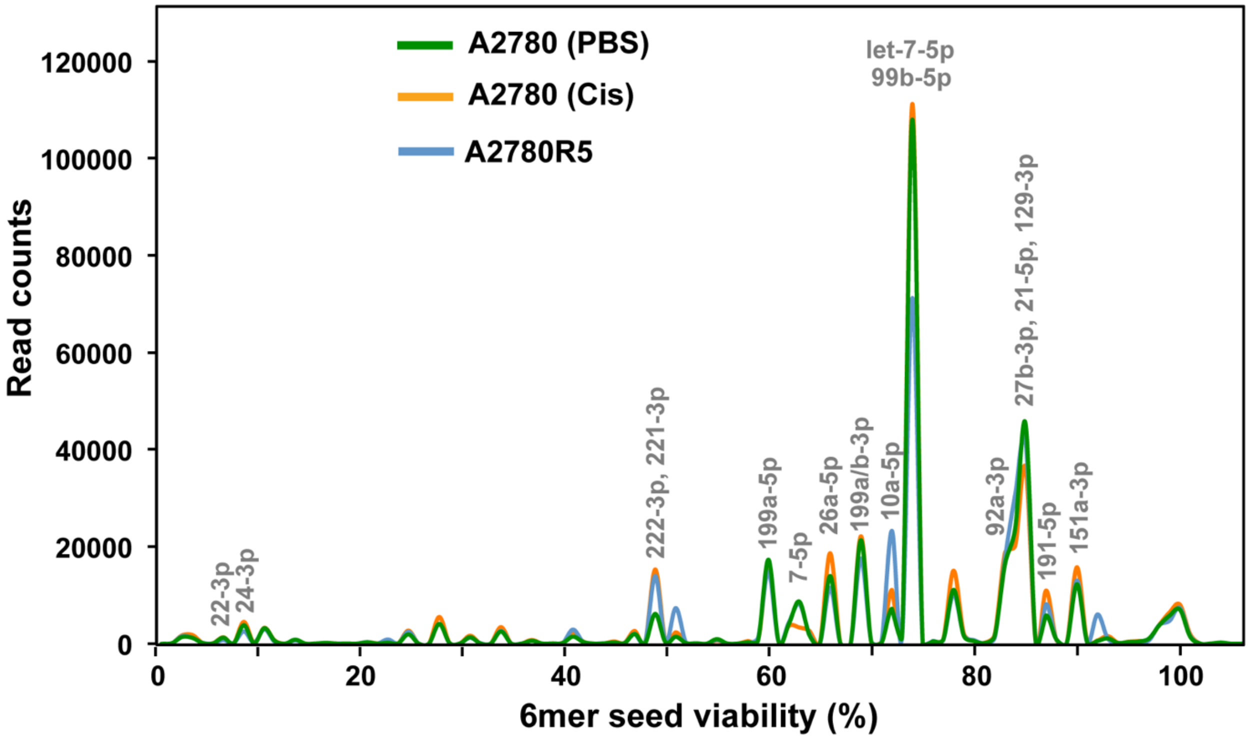


**Supplemental Figure S5.** Effect of cisplatin treatment on Seed Tox composition of the total miRNA fraction. Seed Tox plot of all miRNAs in A2780 cells treated with PBS or cisplatin (Cis) and in A2780R cells. miRNAs that contribute to peaks with more than 5000 reads are labeled. For each labeled peak miRNAs are listed in the order of abundance.

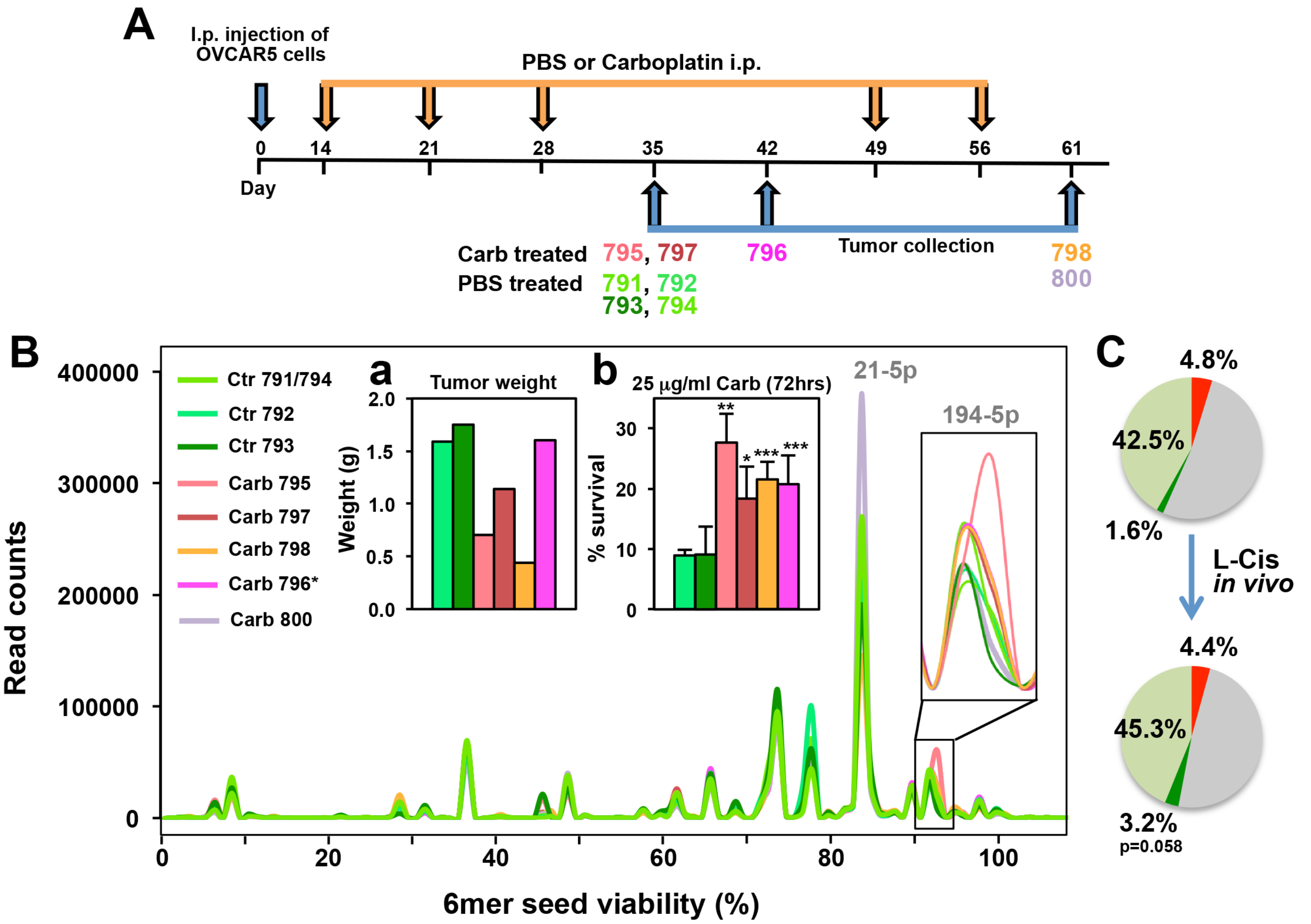


**Supplemental Figure S6**. Effect of long-term *in vivo* carboplatin treatment on Seed Tox of RISC-bound miRNAs in OVCAR5 cells. **A,** Treatment course. OVCAR5 (2 million) cells were injected i.p. into nude mice. Two weeks after inoculation, mice were grouped and treated i.p. with PBS (control, mice ID: 791, 792, 793, 794, n=4), or 25 mg/kg carboplatin (Carb) (n = 5) as indicated: 3 doses of carboplatin (n=2, mice ID: 795, 797); 3-weekly doses of carboplatin + two-week recovery (n = 1, mice ID: 796); 3-weekly doses of carboplatin, followed by 2-weeks recovery, followed by 2-weeks carboplatin (n = 2, mice ID: 798, 800**). B,** Seed Tox plot of total RISC-bound miRNAs in OVCAR5 tumors isolated at different times after PBS or carboplatin treatment as indicated in A. a, Total tumor weight; b, Viability assay of tumors 24 hours after isolated from mice treated as shown in A. ATP assay was performed 96 hrs after treatment. Student's t-test was performed. Significance shown (Student's t-test) is between a carboplatinum treated tumor and either of the two control treated tumors. *** $p<0.0001$, ** $p<0.001$, * $p<0.05$. The area around the miR-194-5p peak is magnified. **C,** Pie charts: Average Seed Tox composition of RISC-bound miRNAs in mice treated with PBS (top) or with carboplatin (bottom). miRNA reads with a predicted 6mer seed viability of <20% are shown in red, >80% in green and 20-80% in grey. Reads of miR-194-5p (part of the green section) are highlighted in darker green. miRNA content of this analysis was 96.1% (average of PBS treated) and 92.6% (carboplatin treated).

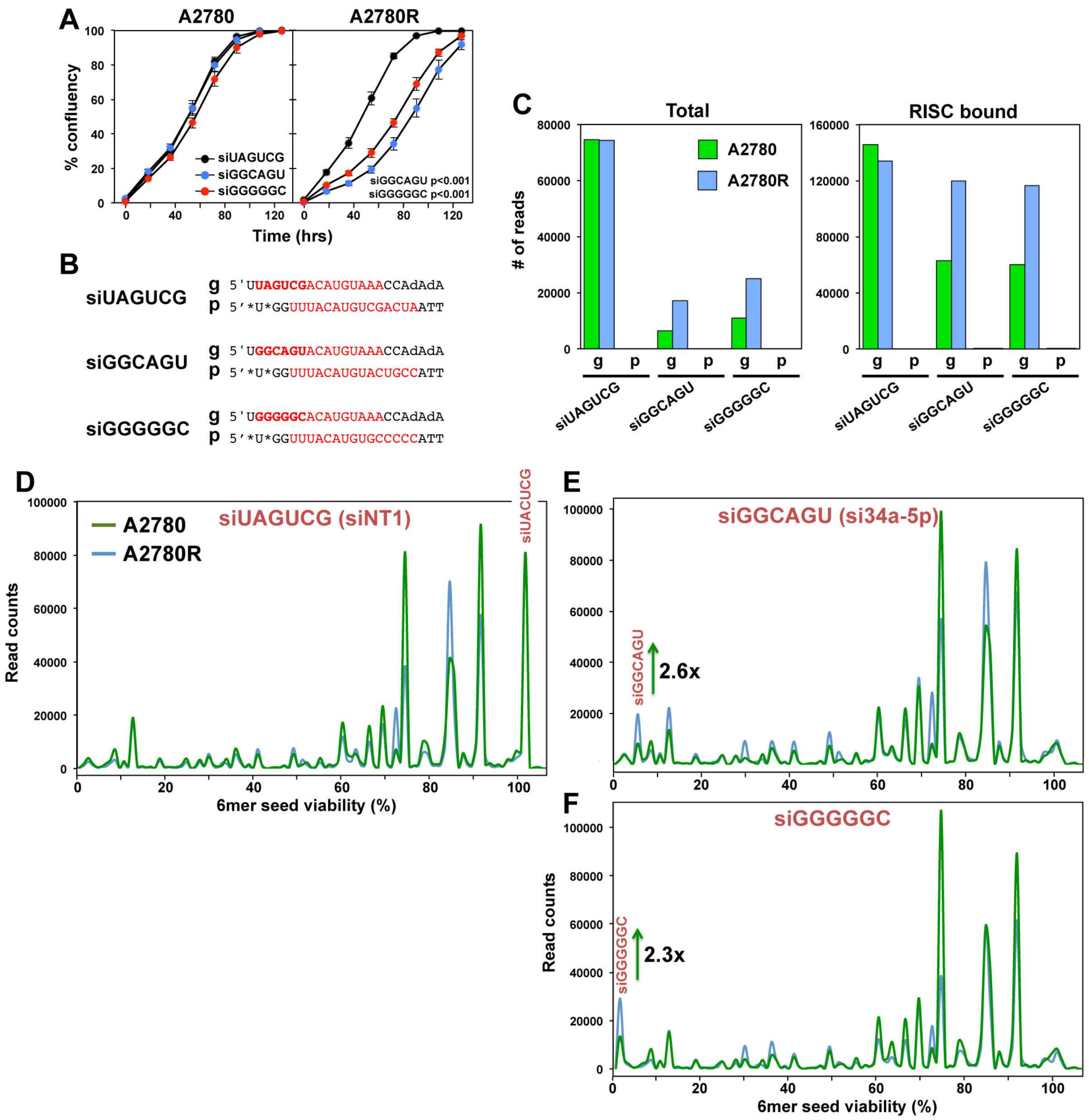


**Supplemental Figure S7.** Preferential uptake of siRNAs with toxic seeds by Pt-R cells. **A,** Change in confluency of A2780/A2780R cells transfected with 1 nM of an siRNA backbone used in the 4096 siRNA screen containing three different 6mer seeds shown in B. P-values were calculated using ANOVA. **B,** Sequences of the three siRNAs differing in only the 6mer seed (6mers bolded). Both guide (g) and passenger (p) strands are shown 5' -> 3'. The sequences used to search the RNA Seq data (as DNA sequences) for presence of exogenous RNAs is shown in red. **C,** Total normalized numbers of sequences derived from the transfected siRNAs [either guide (g) or passenger (p) strand] identified in the total or RISC-bound sRNA in cells transfected with the indicated siRNA. **D-F,** Seed Tox plot of miRNAs and exogenous siRNAs in total smRNA in A2780/A2780R cells 24 hrs after transfection with either siUAGUCG (siNT1), and the highly toxic siGGCAGU or siGGGGGC.

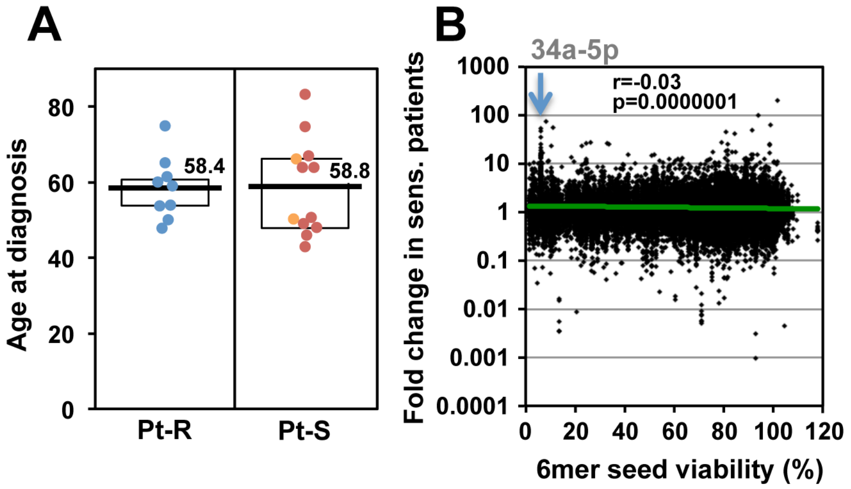


**Supplemental Figure S8.** Lower Seed Tox of RISC-bound sRNAs in OC tumors is associated with increased resistance to chemotherapy. **A,** Differences in age at diagnosis between the two groups in Fig. 4A. **B,** Pearson correlation between the fold change in Pt-S versus Pt-R patients and the 6mer Seed Tox (% viability in HeyA8 cells). Shown are all ~38,000 different reads across all samples. The position of a number of reads representing miR-34a-5p highly enriched in the Pt-S cells is indicated. Note, these reads while highly upregulated, did not reach statistical significance in the fold-change analyses.

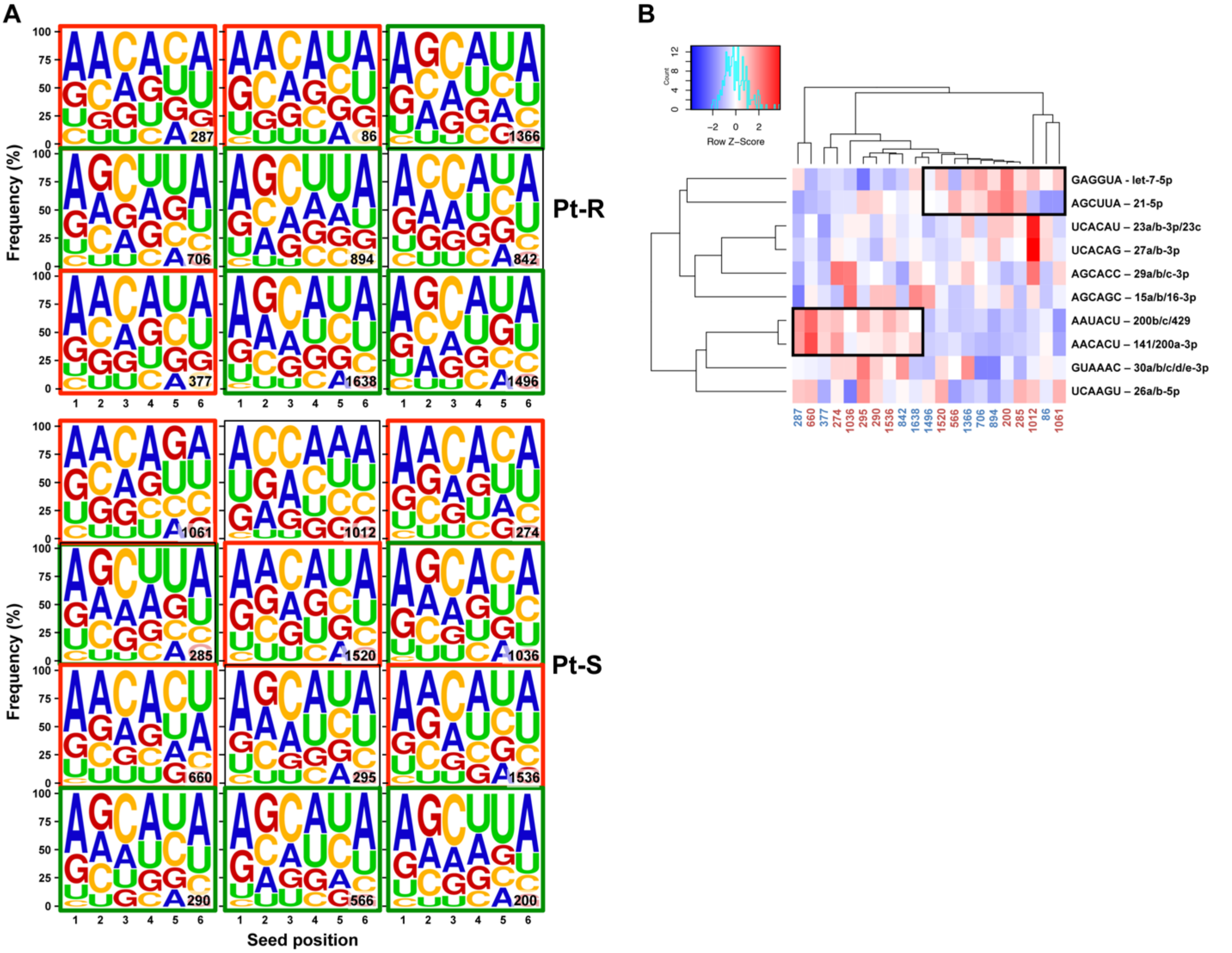


**Supplemental Figure S9.** OC patients can be divided into two subgroups. **A,** Seed composition plots of all RISC-bound sRNAs in all Pt-R and Pt-S tumors. Seeds that are dominated by AAC in the first three positions are boxed in red, the ones dominated by AGC are boxed in green. **B,** Heat map of an unsupervised hierarchical cluster analysis of the ten most abundant RISC-bound seeds in reads in tumors from Pt-S (labeled in blue) and Pt-R (labeled in red) patients. For each 6mer seed, the major miRNA or miRNA families they are part of, are shown. Two clusters of let-7-5p/21-5p and miR-200-5p members, respectively, are boxed.

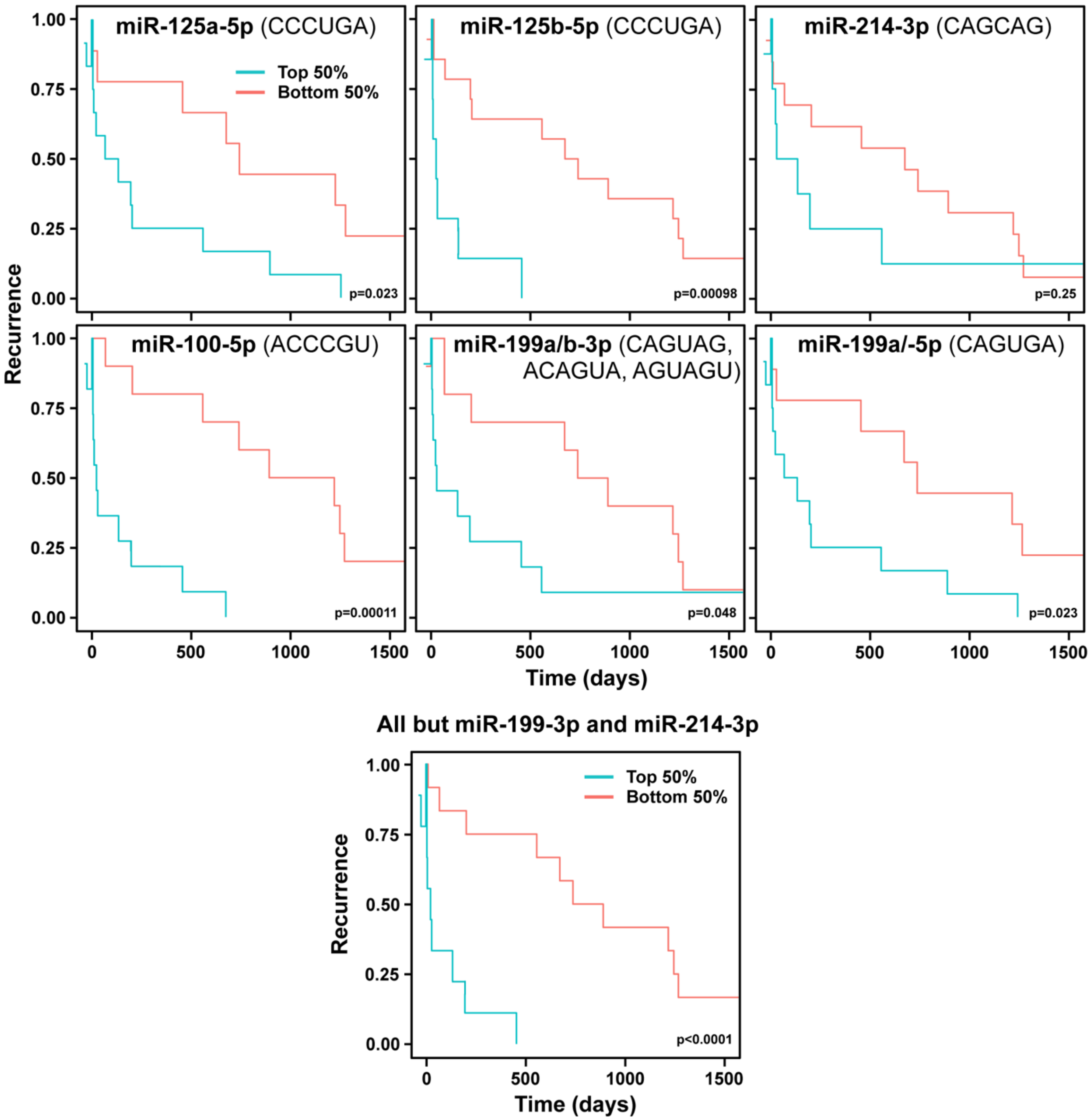


**Supplemental Figure S10.** Kaplan-Meier analyses of individual miRNAs. Kaplan-Meier analysis of patients with the top and bottom highest RISC content of the indicated miRNA. For miR-199-3p the average of all three isomiRs was used.

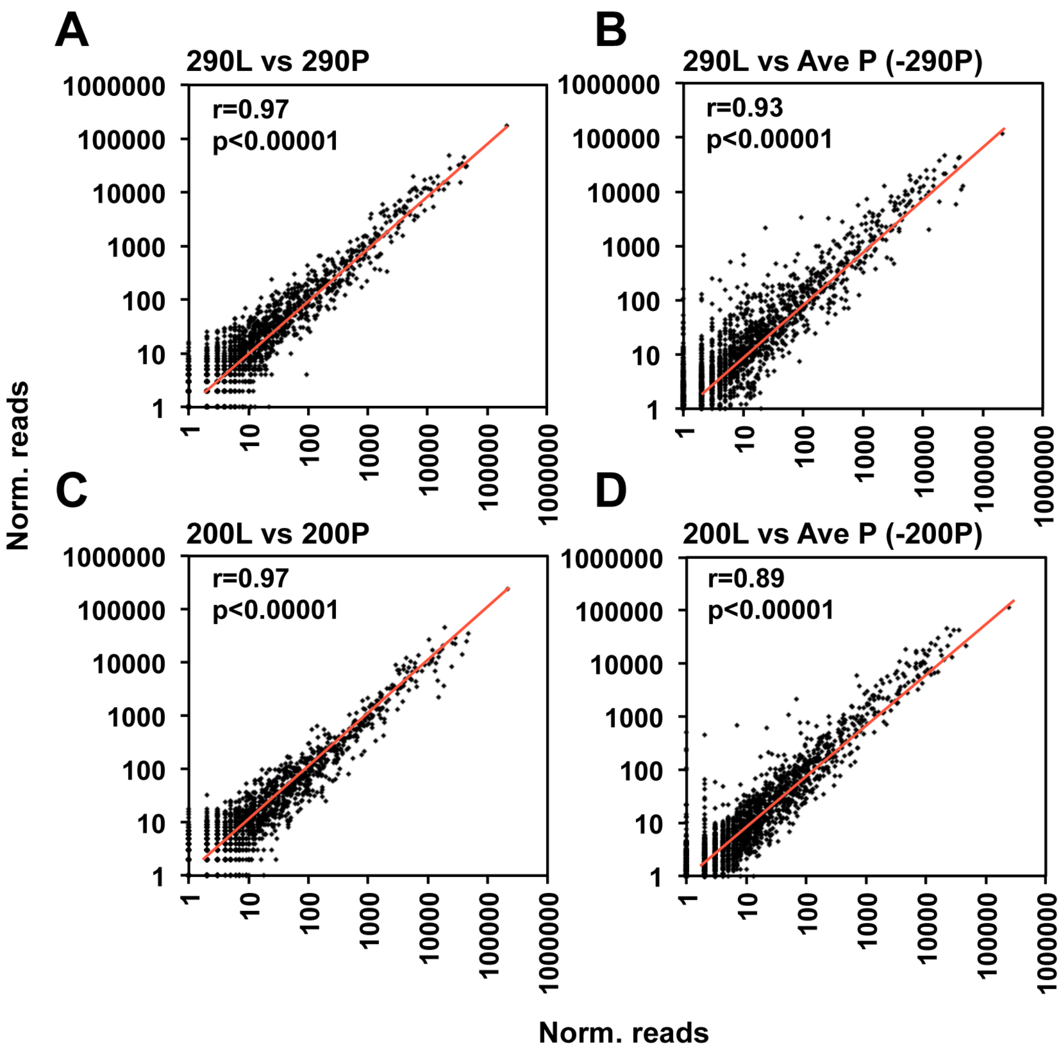


**Supplemental Figure S11.** Reproducibility of the Ago pull down/smRNA Seq analysis in two patients across the two analyses. **A,** Pearson correlation between reads from an Ago pull down analysis of the primary tumor from patient #290 in experiment #1 (short versus long-term survivors) and #2 (primary versus recurrence in long-term survivors). **B,** Pearson correlation between reads from an Ago pull down analysis of patient #290L in experiment #1 versus the average of all patients of experiment #2 minus patient #290P reads. **C,** Pearson correlation between reads from an Ago pull down analysis of the primary tumor from patient #200 in experiment #1 and #2**. D,** Pearson correlation between reads from an Ago pull down analysis of patient #200L in experiment #1 versus the average of all patients of experiment #2 minus patient #200P reads. In all cases all reads were analyzed but only the ones were plotted with an average minimum read count of 1 across all samples. Note, in both cases while the tumors analyzed were from the same patient, the analysis was performed with a different tumor from the same patient.

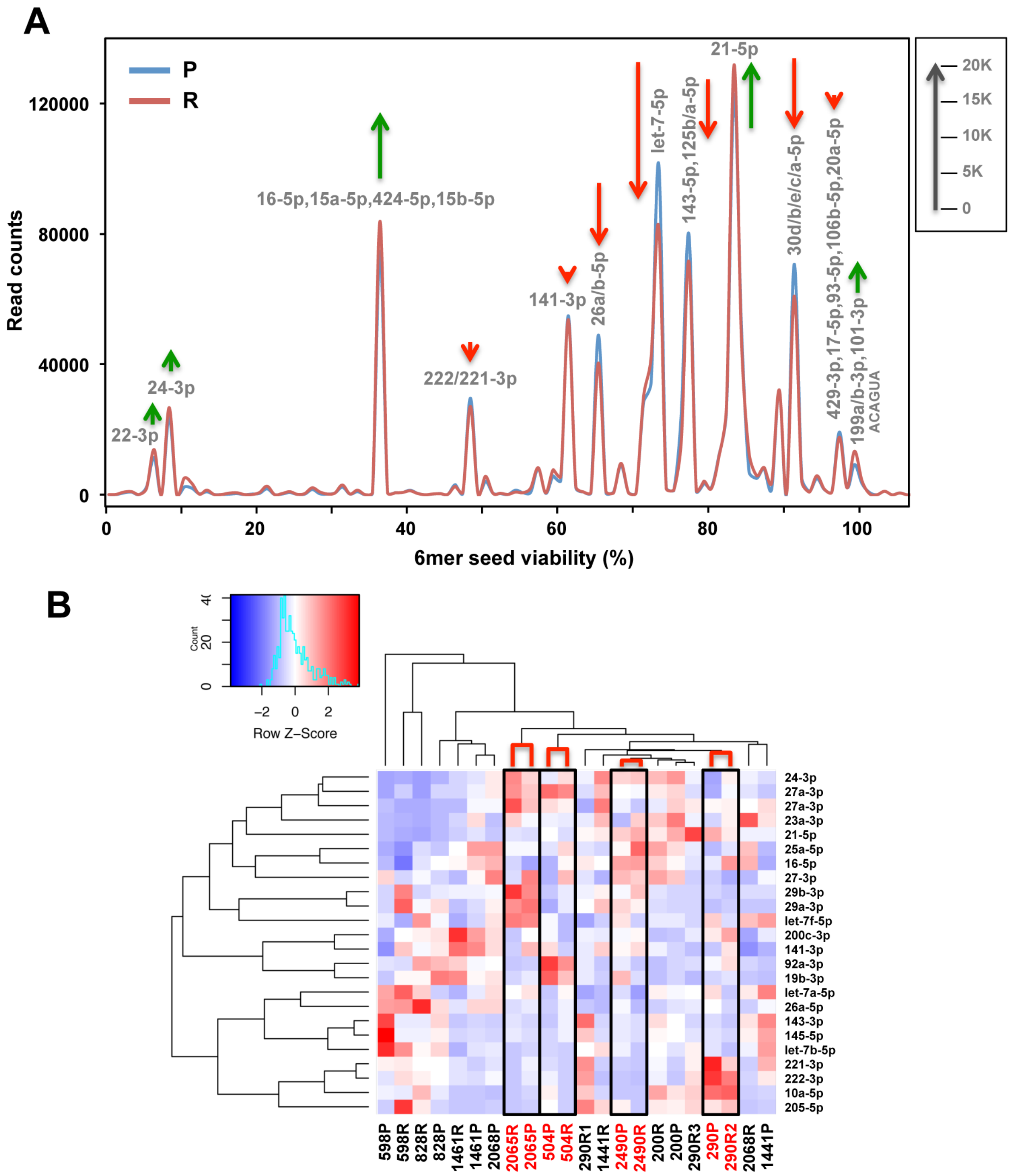


**Supplemental Figure S12.** Analysis of primary tumors and matched recurrences. **A,** Seed Tox plot of all RISC-bound reads in primary tumors and matching recurrences. Green arrows, net increase, and red arrows, net decrease of >1000 reads in peak in recurrent tumors, respectively. **B,** Heat map of an unsupervised hierarchical cluster analysis of the most abundant miRNAs (read average across all samples >10,000) in primary tumors and matching recurrences from 10 patients. Primary tumor and matching recurrence that clustered directly together are boxed. Note, for patient #290 only recurrence R2 clustered together with the primary tumor and this pair was therefore used for further analysis. Patients in a subgroup of four highly related primary/recurrent tumor pairs are shown in red.

### Supplemental Tables

#### The balance between toxic versus nontoxic microRNAs determines platinum sensitivity in ovarian cancer

Monal Patel, Yinu Wang, Elizabeth T. Bartom, Rohin Dhir, Kenneth P. Nephew, Daniela Matei, Andrea E. Murmann, Ernst Lengyel, and Marcus E. Peter

**Table S1: miRNAs involved in therapy resistance or associated with negative treatment outcome in OC**

| Main miRNA | Other miRNAs | Treatment | OvCa cell lines | Type of OvCa-patient sample | Clinical data/relevance | Identified miRNA target | Ref. |
|---|---|---|---|---|---|---|---|
| miR-17~92 cluster | - | Taxol | SKOV3 and Taxol-resistant SKOV3-TR30 | - | - | BMI | 1 |
| miR-17-5p | - | Taxol | OVCAR3, SKOV3 | - | - | PTEN | 93 |
| miR-21 | - | Cisplatin | A2780R (Cis resistant) | - | - | - | 2 |
| miR-21-3p | >20 other miRNAs | Cisplatin | A2780, A2780-CP70, OVCAR5, OVCAR8 and IGROV1 | Publically available datasets | Yes | NAV3 | 3 |
| miR-21-5p | - | Taxol | A2780, PEA1, PEA2, ALST, OVCAR432, OVCAR433, HeyA8, HeyA8-MDR, SKOV3ip, SKOV3-TR, SKOV3, cancer asssociated fibroblasts and adipocytes | OvCa | Yes | APAF1 | 4 |
| miR-21-5p | - | Cisplatin | A2780, A2780- Cp resistant, SKOV-3 | Recurrent patients enrolled in TCGA | Yes | PDCD4 | 5 |
| miR-21-5p | 320 other miRNAs | Cisplatin | SKOV3ip1, HeyA8, A2780, A2780CP20 | - | - | PDCD4 | 6 |
| miR-27a | - | Taxol | A2780 and A2780/Taxol resistant cells | - | - | HIPK2 | 7 |
| miR-31 | - | Cisplatin | OVCAR5, A2780 and its cisplatin-resistant derivatives, MCP1, CP70 | Publically available datasets | Yes | KCNMA1 | 8 |
| miR-93-5p | 7 other miRNAs | Cisplatin | OVCAR3, SKOV3, OVCAR3/CDDP, SKOV3/CDDP | OvCa | Yes | PTEN | 94 |
| miR-98-5p | let-7c-5p, let-7e-5p, let-7g-5p | Cisplatin | C13*, OV2008, A2780, HO8910, SKOV3, CaOV3, Hey, COV362, | Epithelial OvCa | Yes | Dicer1 | 9 |
| miR-106a-5p | - | Cisplatin | OVCAR3, OVCAR3/CIS | - | - | PDCD4 | 95 |
| miR-106a-5p | miR-96, miR-629 | Taxol | SKOV3 and PTX resistant SKOV3 sublines | Ovarian serous tumors | Yes | BCL10, CASP-7 | 10 |
| miR-125b-5p | - | Cisplatin | OV2008 and its resistant variant (C13*) | - | - | Bak1 | 11 |
| miR-125b-2-3p | miR-126-5p | Cisplatin | A2780, A2780CIS, A2780TC1, A2780TC3 | - | - | - | 15 |
| miR-129-5p | - | Taxol | SKOV3, SKOV3/PTX, HeyA8, HeyA8/PTX | - | - | ABCB | 12 |
| miR-130a | miR-374a, miR-146a, miR-182 and 32 others | Cisplatin | A2780, A2780/DDP | - | - | MDR-1/P-gp/PTEN | 13 |
| miR-130a | 54 other miRNAs | Cisplatin | SKOV3, SKOV3/CIS | - | - | MDR1/P-gp | 14 |
| miR-130b | - | Cisplatin, Taxol | A2780, A2780/Taxol | - | - | - | 96 |
| miR-141-3p | 5 other miRNAs | Cisplatin | A2780 and A2780 DDP, TOV112D, TOV21G, OV56, OAW42 | Non-serous tumors | Yes | KEAP1 | 16 |
| miR-149-5p | - | Cisplatin | TOV-21G, A2780, OVCAR-3, Caov-3, ES-2, HO-8910, SK-OV-3 | Ovarian serous cystadenocarcinoma, benign ovarian lesions, TCGA dataset | Yes | Hippo signaling pathway | 18 |
| miR-197-3p | - | Taxol | A2780, SKOV3, A2780/Tax | - | - | NLK | 20 |
| miR-200c-with cytosolic HuR | | Cisplatin, Taxol, Patupilone | - | OvCa patients | Yes | - | 113 |
| miR-214 | miR-100, miR-199a-3p, miR-200a | Cisplatin | A2780S, A2780CP, OV119 | Primary OvCa | Yes | PTEN | 97 |
| miR-214-3p | - | Cisplatin | A2780, A2780cp, OVCAR-3, SKOV3 | - | - | MEG3 | 21 |
| miR-216a-5p | - | Cisplatin | SKOV3, SKOV3 CR, OVCA433 | - | - | PTEN | 22 |
| miR-224-5p | - | Cisplatin | A2780CP/A2780S, C13/OV2008 | Ovarian papillary serous carcinoma | Yes | PRKCD | 23 |
| miR-363-3p | - | Taxol | SKOV3, KF, KF-TX, OVTOKO | OvCa | Yes | LATS2 | 24 |
| miR-376c-3p | - | Cisplatin, Carboplatin | OV2008, A2780 | Serous OvCa | Yes | ALK7 | 98 |
| miR-433 | - | Taxol | A2780, PEO1, PEO4 | - | - | CDK6 | 25 |
| miR-483-3p | - | Cisplatin, Oxiplatin | IGROV-1, IGROV-1/Pt1, IGROV-1/OHP, A2780, A2780/CP, OVCAR5, OVCAR5/Pt | GEO datasets | Yes | PRKCA | 99 |
| miR-490-3p | - | Taxol | A2780, A2780/Taxol | - | - | - | 100 |
| miR-493-3p | - | Taxol | CAOV- 3, OVCAR-8 | High grade serous OvCa | Yes | Mad2 | 26 |
| miR-493-5p | 7 other miRNAs | PARPi, Platinum | BRCA2 mutant KURAMOCHI, OVSAHO, and VU423 cells | BrCa2 mutated OvCa, TCGA cohort | Yes | RNASEH2A, FEN1, SSRP1 | 27 |
| miR-520g-3p | - | Cisplatin, Taxol | A2780, SKOV3, OVA433, ES-2, OV2008, CaOV-3, MCAS, OVK17 | Epithelial OvCa | Yes | DAPK2 | 117 |
| miR-551b-3p | - | Cisplatin | Side population from primary ascites-derived OvCA cells, SKOV3, 8910 | OvCa | Yes | Foxo3, TRIM31 | 28 |
| miR-622 | - | Taxol, Carboplatin | - | High grade serous OvCa | Yes | - | 89 |
| miR-622 | - | Cisplatin, Carboplatin, Olarparib, Veliparib | UWB1.289 | OvCa datasets | Yes | Ku Complex | 101 |
| miR-630 | - | Taxol | SKOV3 and SKOV3-TR | - | - | APAF-1 | 29 |
| miR-1307 | - | Cisplatin, Taxol | SKOV3 and SKOV3- TR30 | Ovarian serous cystadenocarcinoma | Yes | DAPK3 | 32 |
| miR-1307 | - | Taxol | SKOV3 , A2780, A2780/Taxol | - | - | ING5 | 33 |
| let-7e | - | Taxol | A2780, A2780TAX | - | - | - | 15 |

**Supplemental Table S1:** List of miRNAs shown to be involved in therapy resistance or associated with negative treatment outcome in OC. Each row consists of a main miRNA and other miRNAs studied in a published article as well as other details from the study.

**Table S2: miRNAs that can sensitize OvCa cells to therapy or are associated with positive treatment outcome in OC**

| Main miRNA | Other miRNAs | Treatment | OvCa cell lines | Type of OvCa-patient sample | Clinical data/relevance | Identified miRNA target | Ref. |
|---|---|---|---|---|---|---|---|
| miR-7-5p | - | Taxol | HO8910pm | - | - | EGFR, ERK pathway | 43 |
| miR-9 | miR-145, miR-429, miR-26a | Cisplatin | Primary ovarian tumor cells | Serous epithelial OvCa | Yes | - | 44 |
| miR-9-5p | - | Cisplatin, PARP1 inhibitor | OV2008, C13*, SKOV3, A2780, CaOV3 | Serous OvCa | Yes | BRCA1 | 45 |
| miR-15a-5p, miR-16-5p | - | Cisplatin | A2780, A2780- CP20, OVSAHO, OVCAR4, OSE tsT/hTERT | High grade serous OvCa | Yes | BMI1 | 46 |
| miR-18a-3p | miR-15a-5p, miR-25-3p | Docetaxel | SKOV3IP1, HeyA8, HeyA8-MDR, A2780, A2780CP20 | High grade serous OvCa | Yes | K-RAS | 42 |
| miR-29 | - | Cisplatin | CP-70, A2780, HeyAC2, SKOV-3 | - | - | COL1A1 | 37 |
| miR-29b | - | Taxol, Platinum, Cyclophosphamide | OVCAR3 | High grade serous, clear cell, and mucinous ovarian adenocarcinomas | Yes | MAPK10, ATG9A | 38 |
| miR-29b | miR-7, miR-18a, miR-19a | Taxol | ES2, AMOC2 | Epithelial OvCa | Yes | Mcl-1 | 39 |
| miR-30a/c-5p | - | Cisplatin | A2780, A2780/CP70 | - | - | DNMT1, Snail | 103 |
| miR-31 | - | Taxol, Carboplatin | KFrHuman KF OvCa cells and KFr13 cisplatin-resistant KF13, SK-OV-3, OVCAR-3, TU-OM-144 | Serous OvCa | Yes | MET | 40 |
| miR-34c-5p | - | Carboplatin, Docetaxel | OVS1, SKOV-I6 | Ovarian serous cystadenocarcinoma | Yes | AREG | 41 |
| miR-100 | - | Cisplatin | SKOV3, SKOV3/DDP | - | - | mTOR, PLK1 | 47 |
| miR-101-3p | - | Cisplatin | A2780,A2780/DDP, SKOV3, SKOV3/DDP | Epithelial OvCa | Yes | EZH2 | 48 |
| miR-106a-5p | - | Cisplatin | A2780,A2780/DDP | - | - | MCL1 | 49 |
| miR-125b-2-3p | - | Taxol | A2780, A2780TAX | - | - | - | 15 |
| miR-128 | - | Cisplatin | SKOV3, SKOV3/CP | - | - | ABCC5, BMI1 | 50 |
| miR-130a-3p | - | Cisplatin | A2780, A2780/DDP | - | - | XIAP | 104 |
| miR-130a-3p | let-7e, miR-335 | Cisplatin | A2780, A2780CIS, A2780TC1, A2780TC3 | - | - | M-CSF | 15 |
| miR-130b-3p | - | Taxol, Cisplatin | A2780, A2780/CP, A2780/TAX, SKOV3, SKOV3/ TAX | Malignant and benign OvCa | Yes | CSF-1 | 51 |
| miR-133b-3p | - | Taxol, Cisplatin | A2780, A2780/DDP, A2780/Taxol, OVCAR3 | OvCa | Yes | GST-π, MDR1 | 52 |
| miR-134 cluster | - | Taxol | SKOV3, Taxol-resistant SKOV3-TR30 | - | - | c-Myc | 1 |
| miR-134-5p | - | Taxol | SKOV3, Taxol-resistant SKOV3-TR30 | Serous epithelial OvCa | Yes | TAB1 | 53 |
| miR-134-5p | - | Taxol | SKOV3, Taxol-resistant SKOV3-TR30 | Serous epithelial OvCa | Yes | Pak2 | 54 |
| miR-136-5p | - | Taxol | SKOV3 & its PTX-resistant sublines | High grade serous OvCa | Yes | Notch3 | 105 |
| miR-136 | - | Cisplatin | OV2008, C13 | Epithelial OvCa | Yes | - | 55 |
| miR-137 | - | Cisplatin | PEO1, PEO4 , IGROV1, OV90, IGROV1 CR, OV90 CR | OvCa | Yes | EZH2 | 56 |
| miR-139-5p | - | Cisplatin | CAOV3, SNU119, CAOV3/cDDP, SNU119/cDDP | OvCa | Yes | ATP7A/B | 57 |
| miR-139-5p | - | Cisplatin | SKOV3, A2780, SKOV3-R, A2780-R | - | - | c-Jun | 58 |
| miR-141, miR-200a | - | Taxol | SKOV3 | OvCa patient, publicly available datasets | Yes | - | 116 |
| miR-142-5p | - | Cisplatin | OVCAR3, SKOV3 | Epithelial OvCa | Yes | XIAP1, BIRC3, BCL2, BCL2L2, MCL1 | 59 |
| miR-145-5p | - | Taxol | A2780, SKOV3, A2780/PTX, SKOV3/PTX | Serous epithelial OvCa | Yes | Sp1, Cdk6 , P-gp, pRb | 17 |
| miR-146a-5p | - | Cisplatin | OVCAR3, SKOV3 | Primary epithelialOvCa cells from freshly collected malignant ascites | Yes | XIAP, BCL2L2 and BIRC5 | 60 |
| miR-146b-5p | - | Taxol, Cisplatin | SKOV3, HO8910, A2780, OVCAR-3 | Epithelial OvCa | Yes | FBXL10 | 61 |
| miR-149-5p | - | Taxol | A2780 | - | - | MyD88 | 19 |
| miR-152 -3p | - | Cisplatin | C13*, OV2008, A2780, HO8910, SKOV3, CaOV3, Hey, COV362, | Epithelial OvCa | Yes | - | 9 |
| miR-152-3p, miR-185-5p | - | Cisplatin | SKOV3, A2780, A2780/DDP, SKOV3/DDP | - | - | DNMT1 | 62 |
| miR-155-5p | - | Cisplatin | SKOV3, A2780 | Primary epithelial OvCa cells from freshly collected malignant ascites | Yes | XIAP | 63 |
| miR-182 | - | Platinum | - | High grade serous OvCa | Yes | | 64 |
| miR-186-5p | - | Cisplatin | A2780, OV2008, OVCAR3, SKOV3, CAOV3, the related cDDP-resistant cell lines (ACRP, C13* and OVCAR3/ DDP) | Serous OvCa | Yes | Twist1 | 65 |
| miR-186-5p | - | Taxol, Cisplatin | OVCAR3, A2780, A2780/DDP, A2780/Taxol | - | - | ABCB1 | 66 |

| Table S2, cont.: miRNAs that can sensitize OvCa cells to therapy or are associated with positive treatment outcome in OC | | | | | | | |
|---|---|---|---|---|---|---|---|
| Main miRNA | Other miRNAs | Treatment | OvCa cell lines | Type of OvCa-patient sample | Clinical data/relevance | Identified miRNA target | Ref. |
| miR-199a | - | Taxol, Cisplatin, Adriamycin | CD44+/CD117+ ovarian tumor cells from inpatient clinic patients | Epithelial OvCa | Yes | CD44 | 67 |
| miR-199a-3p | - | Cisplatin | SKOV3 , SKOV3-CDDP | OvCa | Yes | ITGB8 | 68 |
| miR-199-3p | - | Cisplatin, 5Aza-dC | SKOV3, HO-8910, IOSE386 (immortalized ovarian epithelial cell line) | Epithelial OvCa | Yes | DDR1 | 106 |
| miR-199b-5p | - | Cisplatin | A2780s, A2780cp, OV2008, C13*, SKOV3, OVCA433, ES-2 | Epithelial OvCa | Yes | JAG1 | 69 |
| miR-200 family | - | Taxol | - | OvCa | Yes | β-tubulin III | 114 |
| miR-200a | - | Taxol | OVCAR3 | - | - | - | 115 |
| miR-200b, miR-200c | - | Cisplatin | HIOSE-80, MCC-3, SKOV3, A2780CP, A2780, OV119 | Primary ovarian tumor | Yes | DNMTs | 70 |
| miR-200c | - | Taxol | HEY, SKOV3, OVCA 420, OV 1847, OVCA 433 | Serous OvCa | Yes | TUBB3 | 71 |
| miR-200c-3p | - | Cisplatin, Olaparib | UWB1.289, UWB1.289-BRCA and SKOV3 | Serous and mucinous carcinomas | Yes | Neuropilin 1 | 72 |
| miR-200c-3p | - | Cisplatin, Taxol, Doxorubicin, Mitomycin C, Vincristine, Epothilone | 2008, Hey, SKOV3, OVCA 420, OVCA 433 | - | - | TUBB3 | 107 |
| miR-200c-with nuclear HuR | | Cisplatin, Taxol, Patupilone | - | OvCa patients | Yes | - | 113 |
| miR-204-5p | - | Cisplatin | A2780, SKOV3, OVCAR3, OV2008, C13* | Epithelial OvCa | Yes | IL-6R | 73 |
| miR-215 | - | Taxol | OVCAR3, CAOV3, SKOV3, HEY | Epithelial OvCa | Yes | XIAP | 74 |
| miR-216b-5p | | Cisplatin | SKOV3, SKOV3/CDDP | OvCa | Yes | PARP1 | 75 |
| miR-335 | - | - | - | Epithelial OvCa | Yes | - | 108 |
| miR-335-5p | 8 other miRNAs | Cisplatin | A2780, A2780/DDP, OV90, OVCAR-3, | - | - | BCL2L2 | 76 |
| miR-338-3p | - | Cisplatin | A2780, A2780/DDP, SKOV3, SKOV3/CDDP | PrimaryOvCa | Yes | WNT2B | 77 |
| miR-363-3p | - | Cisplatin | OV2008, A2780, C13, A2780cp | OvCa | Yes | Snail | 78 |
| miR-370-3p | - | Cisplatin | SKOV3, UWB1.289, HEY, OV2008, IGROV1, TOV112D, ES-2, TOV21G | Endometriod OvCa | Yes | ENG | 109 |
| miR-378a-3p | - | Cisplatin | OVCAR3, SKOV3 | OvCa | Yes | MAPK1, GRB2 | 79 |
| miR-383-5p | - | Taxol | SKOV3, A2780, OVCAR-3, Caov-3 | OvCa | Yes | TRIM27 | 80 |
| miR-411-5p | - | Cisplatin, Taxol | SKOV3, OVCAR3 | Primary OvCa | Yes | ABCG2 | 81 |
| miR-429 | - | Cisplatin | OVCAR3 & Hey | - | - | - | 112 |
| miR-449a | - | Cisplatin | A2780, A2780/CDDP, SKOV3, SKOV3/CDDP | - | - | NOTCH1 | 110 |
| miR-489-3p | - | Cisplatin | SKOV3, OVCAR3, SKOV3/CDDP, OVCAR3/CDDP | - | - | Akt3 | 82 |
| miR-490-3p | - | Cisplatin | SKOV3, OVCAR3, SKOV3/CDDP, OVCAR3/CDDP | OvCa | Yes | ABCC2 | 83 |
| miR-497-5p | 9 other miRNAs | Cisplatin | A2780, A2780/ CP, SKOV3, SKOV3/CP | OvCa | Yes | mTOR, P70S6K1 | 84 |
| miR-503 | - | Cisplatin | SKOV3, SKOV3/DDP | - | - | PI3K p85 | 85 |
| miR-503-5p | - | Taxol | CaOV3, SKOV3, OVCAR3, OV90, CaOV3/PTX-R, SKOV3/PTX-R | - | - | CD97 | 86 |
| miR-506-3p | - | Cisplatin, Olaparib | HeyA8, OVCA433, SKOV3 | Serous OvCa | Yes | RAD-51 | 111 |
| miR-509-3p | - | Cisplatin | SKOV3 | OvCa tissue | Yes | GOLPH3, WLS | 87 |
| miR-514-5p | - | Cisplatin | SKOV3, OVCA433 | NCBI's GEO datasets | Yes | ATP binding cassette subfamily | 88 |
| miR-591 | miR-106a-5p | Taxol | SKOV3 and PTX resistant SKOV3 sublines | Epithelial OvCa | Yes | ZEB1 | 10 |
| miR-708-5p | - | Cisplatin | A2780, A2780/DDP, SKOV3, SKOV3/CDDP | - | | IGF2BP1 | 90 |
| miR-770-5p | - | Cisplatin | OV2008, A2780, C13, A2780cp | Epithelial OvCa | Yes | ERCC2 | 91 |
| miR-873 | - | Cisplatin, Taxol | A2780, A2780/DDP, A2780/Taxol, OVCAR3 | - | - | ABCB1 | 30 |
| miR-874-5p, miR-874-3p | - | Taxol | Caov3, SKOV3 | Epithelial OvCa | Yes | SIK2 | 92 |
| miR-1294 | - | Cisplatin | SKOV3 | Advanced stage OvCa | Yes | IGF1R | 31 |
| let-7d-3p | let-7a | Taxol, Carboplatin | SKOV-3 | High grade serous, endometrioid, mucinous, serous papillary low grade or clear cell OvCa | Yes | - | 34 |
| let-7e-5p | - | Cisplatin | A2780, HO8910, ES2, CAOV3, SKOV3 | Serous epithelial OvCa | Yes | BRCA1, Rad51 | 35 |
| let-7e | - | Cisplatin | A2780, ES2, SKOV3, A2780/CP | - | - | - | 102 |
| let-7g | let-7d | Taxol, Carboplatin, Vinblastine | ADR-RES, OVCAR-8, T47D, IGROV1 | Epithelial OvCa | Yes | MDR1, IMP-1 | 36 |

**Supplemental Table S2:** List of miRNAs shown to either sensitize OC to therapy or associated with positive treatment outcome in OC. Each row consists of a main miRNA and other miRNAs studied in a published article as well as other details from the study.

**Supplemental Table S3:** Patient tumor samples used in analysis #1 (long-term vs. short-term survivors) and analysis #2 (primary vs. recurrent tumors).

**Supplemental Table S4:** Identification of the most abundant miRNAs enriched in Pt-R patients. Tab "All": A list of all reads collapsed into miRNAs according to their 6mer seeds in analysis #1 (comparison of short term and long term Pt sensitive patients). S, short-term sensitive; I, intermediate sensitive; L, long-term sensitive. Tab "Top abundant miRNAs". miRNA read numbers were correlated using Pearson correlations with the Pt sensitive days. All miRNAs as shown that had a correlation p value of <0.05 an average read number of >1000 and a downregulation of >1.5 fold between the Pt-S and Pt-R groups.

**References Supplemental Table S1 and S2**
